# Different forms of autophagy restrict neurite outgrowth in disparate compartments in a single neuron

**DOI:** 10.64898/2026.08.24.746867

**Authors:** Sohitri Mukherjee, Andrea Cuentas-Condori, Daniel A. Colón-Ramos, Andrea K.H. Stavoe

**Affiliations:** Department of Neurobiology and Anatomy, McGovern Medical School, University of Texas Health Science Center at Houston, Houston, TX; Department of Cell Biology and Department of Neuroscience, Yale University, New Haven, CT; Wu Tsai Institute, Yale University, New Haven, CT; Instituto de Neurobiología, Recinto de Ciencias Médicas, Universidad de Puerto Rico, San Juan 00901, Puerto Rico

## Abstract

Autophagy is a degradative pathway that is critical in neurons to maintain their homeostasis and to direct nervous system development. Neurons are large, highly polarized cells with distinct compartments that perform discrete functions. How an individual neuron can differentially mobilize autophagy in the axon, dendrite, and cell body is unknown. Here we interrogate the role of autophagy in the neurodevelopment of a single neuron in *Caenorhabditis elegans* to identify how the spatial compartmentalization of neuronal autophagy ultimately restricts neurite outgrowth in multiple neuronal compartments. However, while canonical autophagy restricts dendrite outgrowth, noncanonical forms of autophagy appear to restrain neurite outgrowth in the axon and soma. Through mutant analysis of the autophagy pathway, we identify that WIPI2-independent autophagy modulates ectopic neurite formation in the soma and that ATG9-independent autophagy regulates axon arborization. Further, we find that unrelated lipid scramblases can compensate for the loss of ATG9 in axon arborization. Our data indicate that neurons marshal both canonical and non-canonical autophagy to spatially control development of separate compartments.

## INTRODUCTION

Bulk macroautophagy (hereafter referred to as autophagy), involves the de novo formation of a double-membrane vesicle that sequesters subcellular cargo for delivery to the lysosome for degradation. Autophagy is a homeostatic process that maintains the cellular environment by quarantines damaged organelles and aggregated proteins prior to recycling. Autophagy is particularly important in maintaining neuronal homeostasis and function because neurons are post-mitotic, highly metabolically active, long-lived, and have complex, polarized architectures. The importance of autophagy in neuronal health is underscored by neuron-specific knock out of autophagy genes in mice causing early neurodegeneration and that misregulation of autophagy has been implicated in age-related neurodegenerative diseases including Alzheimer’s, Parkinson’s, and Huntington’s diseases.

The autophagy pathway centers on the formation of a double-membrane autophagosome. Multiple protein complexes composed of 40+ proteins in mammals coordinate autophagosome biogenesis. The initiation/induction complex contains unc-51-like kinase 1/2 (ULK1/2), autophagy-related gene (ATG) 13 (ATG13), ATG101, and FIP200. ULK1 phosphorylates components of other complexes, and the initiation complex is thought to be a master regulator of autophagosome formation. The nucleation complex is composed of ATG14, BECN1, NRFB2, p150, and VPS34. As a Class III phosphatidylinositol 3-kinase complex, it generates PtdIns(3)P. The elongation complex is itself composed of two ubiquitin-like conjugation complexes: the ATG12 conjugation complex and the ATG8 conjugation complex. ATG7 and ATG10 catalyze the conjugation of ATG12 to ATG5. Along with ATG16L1, ATG12-ATG5 form an E3-like complex that works with ATG7 and ATG3 to conjugate ATG8 family members to phosphatidylethanolamine (PE). ATG4 proteases cleave ATG8 proteins in preparation for conjugation to PE. WIPI1 and WIPI2 proteins bridge the nucleation and elongation complexes by binding to ATG16L1 and to PI3P at the developing autophagosome. The ATG2 complex is composed of ATG2, ATG9, and WIPI4. ATG2 is a bridge-like lipid transfer protein that interacts with ATG9, a lipid scramblase and the only transmembrane protein in the core autophagy machinery, to deliver lipids to both leaflets of the growing autophagosome membrane. Autophagosome closure is then mediated by ESCRT machinery (Melia *et al*., 2020).

Neurons are unique cells with highly polarized architectures and much of their cytoplasmic volumes and plasma membrane surface area exist in the axons and dendrites, away from the buffering capabilities of the cell body. Autophagy is spatially regulated in neurons. In primary mammalian neurons, autophagosome biogenesis occurs primarily in the distal axon. After formation, autophagosomes are transported back to the cell body for cargo degradation and recycling. This compartmentalization of neuronal autophagy has also been confirmed *in vivo* in *Drosophila* and *Caenorhabditis elegans*. However, how a neuron coordinates specific roles for autophagy across compartments during development is less clear.

We were interested to address this gap in knowledge in the nematode *Caenorhabditis elegans*. We previously found that autophagy instructs presynaptic assembly in interneuron AIY and axon outgrowth in sensory neuron PVD (Stavoe *et al*., 2016). However, we only previously interrogated autophagy in a single compartment, the axon. Here, we interrogated how autophagy directs the development of sensory motoneuron NSM and how NSM manages autophagy across its disparate compartments. NSM is the only neuron in the hermaphrodite that has a complex axonal arbor. The NSM cell body is positioned in pharynx, anterior to the nerve ring. A neurite exits the posterior side of the NSM soma and bifurcates into a dendrite and an axon. The axon further bifurcates into a dorsal and a ventral axon (Figure 1A). The ventral axon later develops a complex arbor starting at larval stage 4 (L4) and arborization continues into young adulthood (Axäng *et al*., 2008; Nelson and Colón-Ramos, 2013). Thus, NSM presented a compelling model to assess how neurons control autophagy across complex architectures.

**Figure 1.**
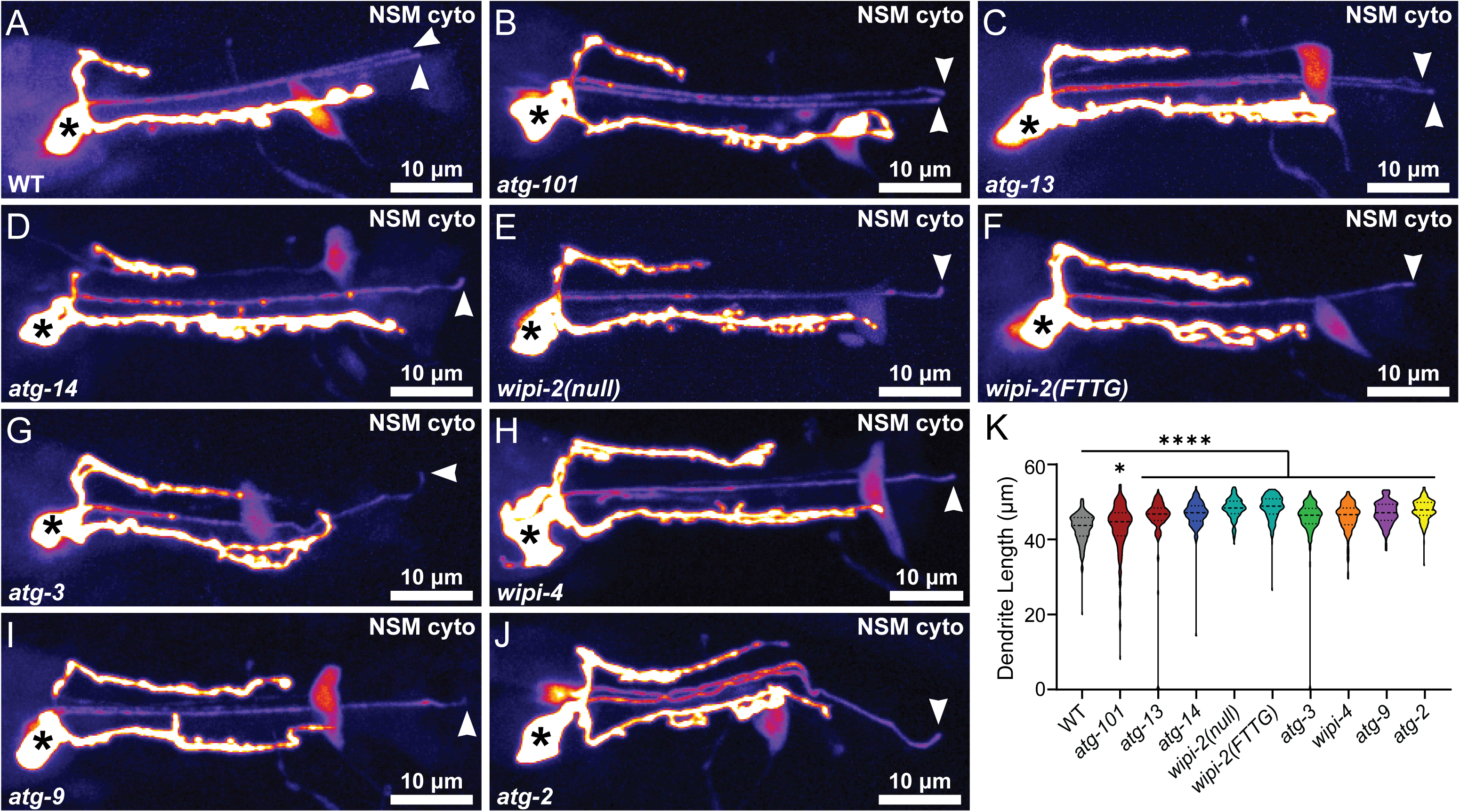
Autophagy components are required to constrain NSM dendrite outgrowth. (A-J) Representative maximal projection micrographs of NSM in L4 worms in wild-type (A), *atg-101/epg-9* (B), *atg-13* (C), *atg-14/epg-8* (D), *wipi-2/atg-18(null)* (E), *wipi-2/atg-18(FTTG)* (F), *atg-3* (G), *wipi-4/epg-6* (H), *atg-9* (I), and *atg-2* (J) mutant worms. Micrographs are displayed with the Fire lookup table to more easily visualize the dim NSM dendrite. The maximal projections occasionally include dendrites from both NSM neurons (A-C). Arrowhead indicates the distal end of the dendrite; asterisk identifies the cell body. (K) Quantification of dendrite length. Truncated violin plot with medium smoothing and median and quartiles displayed (n ≥ 135 cells imaged on at least three independent days for each genotype). ****p < 0.0001; *p < 0.05 between mutants and wild-type (as indicated) by Kruskal-Wallis test with Dunn’s Multiple Comparisons.

In this study, we interrogated how autophagy regulates NSM neurodevelopment in each of its compartments: soma, dendrite, dorsal axon, and ventral axon. We determined that autophagy contributes to the proper development of each compartment, but that different forms of autophagy are necessary for the neurodevelopment of distinct compartments. While canonical autophagy regulates NSM dendrite outgrowth, we identified two novel noncanonical forms of autophagy that modulate the neurodevelopment of other compartments. We found that WIPI2-independent autophagy prevents ectopic neurite formation in the soma. Furthermore, we discovered that ATG9-independent autophagy restrains ventral axon arborization. Intriguingly, we determined that other autophagy-associated lipid scramblases can compensate for loss of *atg-9* in NSM ventral axon arborization. We propose that NSM directs compartment-specific development by spatially implementing different forms of autophagy.

## RESULTS

### Canonical autophagy restricts NSM dendritic outgrowth

To visualize NSM morphology, we used a transgenic strain that expresses cytoplasmic GFP primarily in NSM, with limited expression in other serotonergic neurons HSN and AFD (Nelson and Colón-Ramos, 2013). To assess how autophagy regulates NSM development, we first focused on the NSM dendrite (Fig. 1A-J). While multiple studies have uncovered the role of autophagy in axonal development, very little is known about the function of autophagy in the dendrite. We examined putative null or loss-of-function mutants for constituents of each autophagosome biogenesis complex. We examined null alleles of initiation complex members *atg-13* and *atg-101* and the autophagy-specific constituent of the nucleation complex *atg-14*. We examined two alleles of *wipi-2*: a null allele and “FTTG” which mutates R228-R229 to T228-T229. The FRRG motif is required for WIPI binding to PI3P and WIPI autophagic activity; the FTTG allele abrogates both PI3P binding and autophagy (Dove *et al*., 2004; Proikas-Cezanne *et al*., 2007; Lu *et al*., 2011). We used a hypomorphic allele of *atg-3* to assess the involvement of the elongation complex and examined null alleles of *wipi-4, atg-2,* and *atg-9*, the only transmembrane constituent of the core autophagy machinery (Holzer *et al*., 2024). We will refer to autophagy alleles by their mammalian ortholog names – see Table S1 for ortholog and allele information.

We quantified NSM dendrite length at L4. We found that wild-type L4 animals had a median NSM dendrite length of 43.58 µm (Fig. 1A, 1K). In contrast, all of the autophagy mutants displayed significantly longer NSM dendrite lengths at L4 than wild type (Fig. 1B-K). Occasionally we would detect autophagy mutant animals that lacked an NSM dendrite, which we never observed in wild type animals (length of 0 μm in Fig. 1K). *atg-101* mutant worms (Fig. 1B) had a median NSM dendrite length of 44.59 µm (Fig. 1K), which was at the bottom of the range of dendrite length, but all of the other autophagy mutants clustered together (46.31 – 48.71 µm; Fig. 1C-K)). Since loss of autophagy resulted in longer NSM dendrites, these data suggest that autophagy normally works to restrain dendrite outgrowth in NSM. Further, since the array of autophagy mutants that we examined represent all autophagosome biogenesis complexes, our data suggest that canonical autophagy is required to restrict NSM dendrite outgrowth. While this role for autophagy in neuronal development is novel, it is consistent with our previous findings in axon outgrowth in a different neuron, PVD (Stavoe *et al*., 2016), suggesting that autophagy might broadly constrain neurite outgrowth during neurodevelopment.

We next asked if autophagy was acting cell-autonomously to restrict NSM dendrite outgrowth (Fig. S1A-D). We expressed *atg-13* cDNA under the same NSM-specific promoter (tph-1p) that expresses GFP in the NSM reporter line. We generated two independent lines that express this array in *atg-13* mutant animals. In *atg-13* mutant animals that express the rescuing array in NSM, we found that NSM dendrites were significantly shorter (median lengths of 37.65 and 35.76 µm for rescuing lines 1 and 2, respectively; Fig. S1B-D) than *atg-13* mutants (median length of 46.60 µm; Fig. S1A, S1D). We also found that NSM-rescued worms had dendrites that were significantly shorter than wild-type animals (median length 43.58 µm), suggesting that we over-expressed *atg-13* in NSM and those worms have over-active autophagy in NSM. Together, our data indicated that autophagy act cell-autonomously in NSM to restrict NSM dendrite outgrowth.

### Select autophagy complexes differentially affect NSM dorsal axon outgrowth

We next examined how autophagy influenced the morphology of the NSM dorsal axon (Fig. 2A-J). We measured the length of the dorsal axon from the point of axon bifurcation to the distal end of the dorsal axon. We observed that autophagy mutant worms broadly showed more variability in NSM dorsal axon length than wild type animals (Fig. 2B-K). The median NSM dorsal axon length in wild-type animals was 20.02 µm with a standard deviation of 6.82. In contrast, while several autophagy mutants did not display average dorsal axon lengths that were significantly different from wild-type (Fig. 2K), autophagy mutants did consistently display more variable dorsal axon lengths (standard deviations ranged 9.51 – 12.49). Specifically, we found that *atg-101* and *atg-13* mutant worms (Fig. 2B, 2C) had significantly shorter NSM dorsal axons than wild type worms, with median lengths of 12.76 and 14.46 µm, respectively (Figure 2K). In contrast, *atg-3* and *wipi-4* mutant worms (Fig. 2G, 2H) displayed significantly longer NSM dorsal axons compared to wild type animals, with median lengths of 24.63 and 28.48 µm, respectively. Our data suggest that there is some stochasticity in dorsal axon length in autophagy mutants. The role of autophagic components in NSM dorsal axon development is appreciably different than their role in NSM dendrite outgrowth, implying that NSM spatially deploys autophagy to modulate the development of specific neurites. However, our data also indicate that autophagy components are modulating neurite outgrowth in distinct compartments.

**Figure 2.**
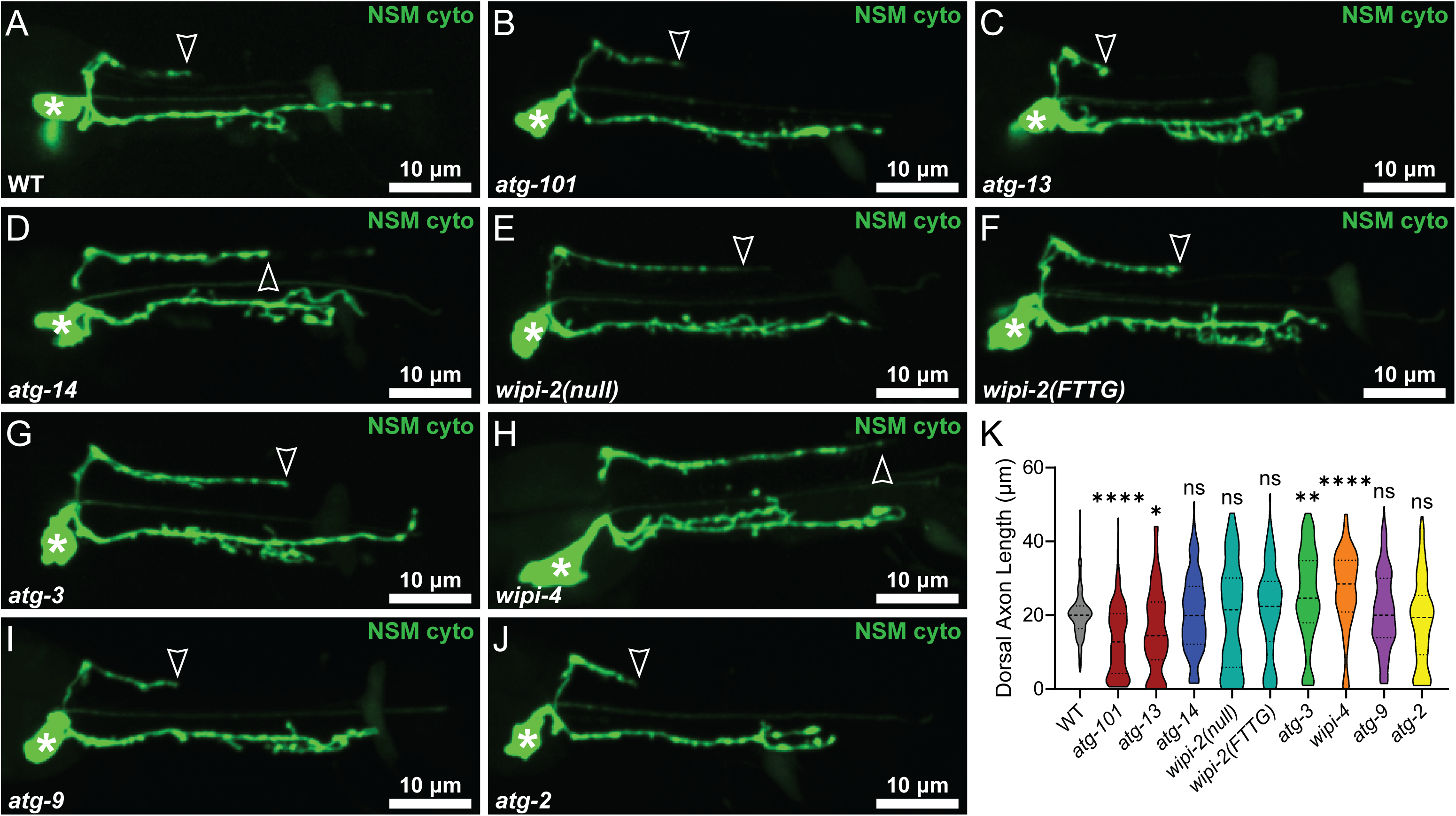
Autophagy components differentially regulate NSM dorsal axon outgrowth. (A-J) Representative maximal projection micrographs of NSM in L4 worms in wild-type (A), *atg-101/epg-9* (B), *atg-13* (C), *atg-14/epg-8* (D), *wipi-2/atg-18(null)* (E), *wipi-2/atg-18(FTTG)* (F), *atg-3* (G), *wipi-4/epg-6* (H), *atg-9* (I), and *atg-2* (J) mutant worms. Open arrowhead indicates the distal end of the dorsal axon; asterisk identifies the cell body. (K) Quantification of dorsal axon length. Truncated violin plot with medium smoothing and median and quartiles displayed (n ≥ 86 cells imaged on at least three independent days for each genotype). ****p < 0.0001; **p < 0.005; *p < 0.05 between wild-type and indicated mutant by Kruskal-Wallis test with Dunn’s Multiple Comparisons.

### WIPI2-independent autophagy restrains NSM ectopic neurite outgrowth

We next examined how autophagy affects the NSM cell body. We observed that autophagy mutants had ectopic neurites emerging from the NSM cell body (Fig. 3A-J). In contrast to the primary process that emerges from the posterior NSM soma, these ectopic neurites frequently emerged from the anterior NSM soma. First, we quantified the fraction of cells that had one or more ectopic neurites. While we detected variability between the autophagy mutants, all autophagy mutants showed a significantly higher frequency of NSM ectopic neurites compared to wild type animals (Fig. 3K). We were intrigued to note that *wipi-2* mutants had the lowest frequency of NSM ectopic neurites among the autophagy mutants. Second, we quantified the number of NSM ectopic neurites per cell. We found that each autophagy mutant examined, with the exception of *wipi-2* mutants, had significantly more ectopic neurites than wild type worms (Fig. 3L).

**Figure 3.**
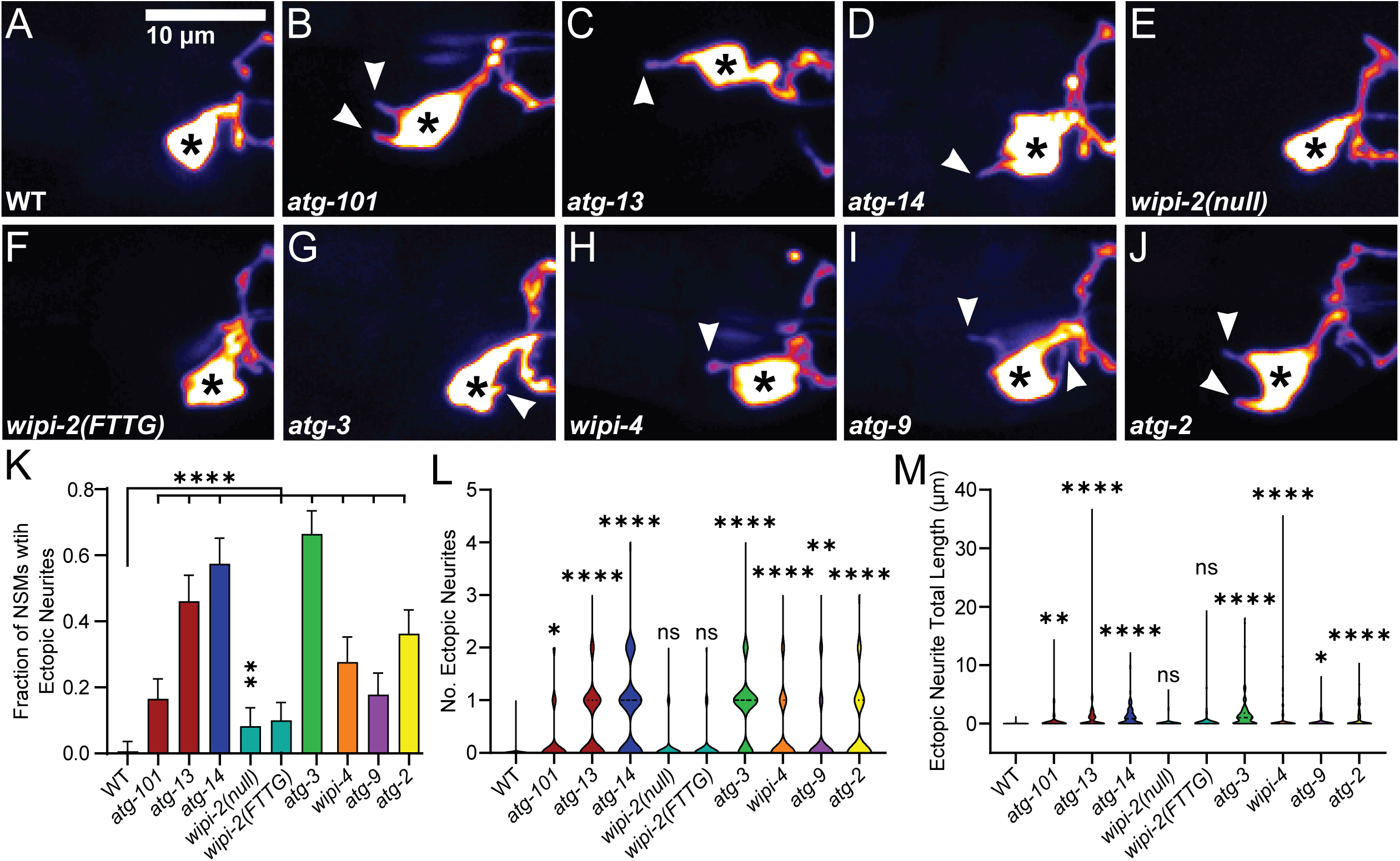
Autophagy components other than WIPI-2 are inhibit ectopic neurites in NSM. (A-J) Representative maximal projection micrographs of NSM in L4 worms in wild-type (A), *atg-101/epg-9* (B), *atg-13* (C), *atg-14/epg-8* (D), *wipi-2/atg-18(null)* (E), *wipi-2/atg-18(FTTG)* (F), *atg-3* (G), *wipi-4/epg-6* (H), *atg-9* (I), and *atg-2* (J) mutant worms. Micrographs are displayed with the Fire lookup table to more easily visualize the dim NSM ectopic neurites. Arrowheads indicate ectopic neurites; asterisk identifies the cell body. (K) Quantification of the penetrance of NSM ectopic neurites in wild-type and autophagy mutant animals. Error bars indicated 95% confidence interval; n > 140 cells imaged on at least three independent days for each genotype. ****p < 0.0001; **p < 0.005 between autophagy mutants and wild-type by Fisher’s exact test. (L-M) Quantification of the number of ectopic neurites per cell (L) and total length of ectopic neurites per cell (M). Truncated violin plot with medium smoothing and median and quartiles displayed (n ≥ 143 cells imaged on at least three independent days for each genotype). ****p < 0.0001; **p < 0.005; *p < 0.05; ns > 0.05 between mutants and wild-type (as indicated) by Kruskal-Wallis test with Dunn’s Multiple Comparisons.

Since we noted that some autophagy mutant animals had very long ectopic neurites, we also quantified the combined length of all ectopic neurites of each NSM neuron. We similarly found that other than *wipi-2* mutants, autophagy mutant animals displayed longer NSM ectopic neurites than wild-type animals (Figure 1L). We were surprised to determine that both *wipi-2(null)* and PI3P-binding deficient *wipi-2(FTTG)* mutant animals did not display significantly more or longer NSM ectopic neurites than wild type animals, particularly since *wipi-4* mutants did phenocopy other autophagy mutants. To our knowledge, WIPI2-independent autophagy that requires ATG-9 and the conjugation complex (represented by ATG-3) has not been previously reported. Together, our data suggest that WIPI2-indpendent autophagy is required for restricting ectopic neurite outgrowth during NSM development.

We next asked whether autophagy acts cell-autonomously to restrain ectopic neurite formation in NSM. As before, we ectopically expressed *atg-13* cDNA using an NSM-specific promoter in *atg-13* mutants, generating two independent rescue lines (Fig. S1A-C). We found that the fraction of NSMs with ectopic neurites (Fig. S1E), the number of NSM ectopic neurites (Fig. S1F), and the total length of ectopic neurites decreased (Fig. S1G) significantly compared to *atg-13* mutants, but were still significantly higher than wild-type worms (Fig. S1A-E). Together, our data suggest that WIPI2-independent autophagy acts cell-autonomously in NSM to prevent ectopic neurite formation, reminiscent of autophagy restricting NSM dendrite outgrowth.

### Most autophagy components are required to restrict NSM ventral axon arborization

We next investigated how autophagy regulates the NSM ventral axon. The NSM ventral axon forms an axonal arbor during development. Beginning at larval stage 4 (L4), branches begin to grow out from the ventral axon in the vicinity of the nerve ring. This arbor continues to develop and increase in complexity into young adulthood (Nelson and Colón-Ramos, 2013). We next asked whether autophagy regulated the development of the NSM axon arbor. We observed that most autophagy mutants displayed more complex axon arbors at L4 compared to wild type animals (Fig. 4A-4J). To quantify the axon arbor complexity, we first isolated the ventral axon from the maximal projection micrograph of a z-stack containing a single NSM cell. We then developed an unbiased analysis pipeline in Nikon Elements using General Analysis 3 (GA3). Using the GA3 pipeline, we quantified the ventral arbor complexity by measuring total ventral axon length (including all branches), the number of branchpoints in the ventral axon, and the fluorescence intensity of the ventral axon.

**Figure 4.**
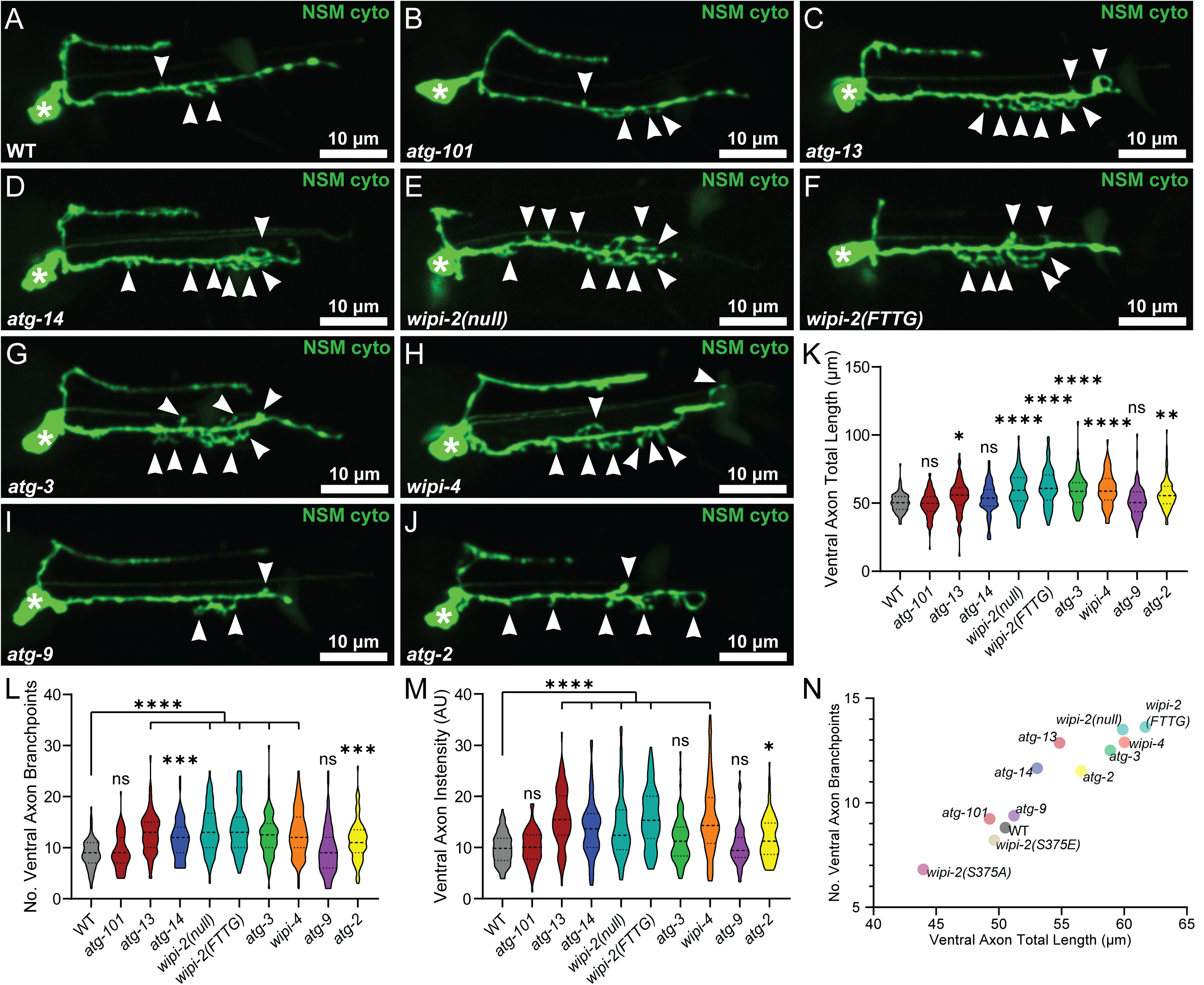
Autophagy components, but not ATG-9, are required to constrain NSM ventral axon arbor complexity. (A-J) Representative maximal projection micrographs of NSM in L4 worms in wild-type (A), *atg-101/epg-9* (B), *atg-13* (C), *atg-14/epg-8* (D), *wipi-2/atg-18(null)* (E), *wipi-2/atg-18(FTTG)* (F), *atg-3* (G), *wipi-4/epg-6* (H), *atg-9* (I), and *atg-2* (J) mutant worms. Arrowheads indicate ventral axon branches; asterisk identifies the cell body. (K-M) Quantification of the total length of the ventral axon arbor, including branches (K), the number of ventral axon branchpoints (L) and the total intensity of the ventral axon arbor (M). Truncated violin plots with medium smoothing and median and quartiles displayed (n ≥ 71 cells imaged on at least three independent days for each genotype). ****p < 0.0001; ***p < 0.0005; **p < 0.005; *p < 0.05; ns > 0.05 between mutants and wild-type by Kruskal-Wallis test with Dunn’s Multiple Comparisons. AU, arbitrary units. (N) Plot of the means for number of branchpoints (y-axis) and ventral axon total length (x-axis) for indicated genotypes.

We found that most autophagy mutants displayed significantly higher ventral axon total length (Fig. 4K), more ventral axon branchpoints (Fig. 4L), and higher ventral axon fluorescence intensity (Fig. 4M) than wild type animals. However, we were surprised to find that *atg-101* and *atg-9* mutant worms did not display significant differences from wild type animals in any of the three measurements (Fig. 4K-4M). We plotted the mean values for all animals for the number of branchpoints versus the axon total length to more easily compare wild type and autophagy mutant animals (Fig. 4N). The means of *atg-101* and *atg-9* mutant worms cluster very close to wild type, in contrast to other autophagy mutants (Fig. 4N). Since all autophagy mutants other than *atg-101* and *atg-9* yielded more complex NSM ventral axon arbors, our data suggest that a non-canonical type of autophagy regulates ventral axon arborization.

We again assessed whether autophagy acts cell-autonomously to regulate ventral axon arborization. Using the same *atg-13* rescuing strategy (Fig. S1A-C), we found that *atg-13* mutant animals from both rescuing lines that ectopically express *atg-13* cDNA in NSM had significantly shorter ventral axon total length and fewer ventral axon branchpoints than *atg-13* mutant animals (Fig. S1H-J). Similar to our results in dendrite length and ectopic neurites, the *atg-13* rescuing lines displayed significantly shorter ventral axon total length and fewer branchpoints than wild-type animals, suggesting that the rescuing lines over-activated autophagy (Fig. S1H-J). We did not assess ventral axon fluorescence because we used the same promoter to drive NSM-specific expression in both the original transgenic strain to visualize NSM morphology and the rescuing array. Together, our data suggest that non-canonical autophagy acts cell-autonomously to restrain NSM ventral axon arborization.

We previously found that phosphorylation of WIPI2 can modulate autophagy in both mouse (WIPI2 Serine 395; (Stavoe *et al*., 2019)) and *C. elegans* neurons (WIPI2/ATG-18 Serine 375; (Tsong *et al*., 2026)), with WIPI2 phospho-dead constructs permitting autophagosome biogenesis to begin and WIPI2 phospho-mimetic constructs inhibiting autophagosome formation. We next asked whether WIPI2 phosphorylation affected the neurodevelopment of any of the NSM compartments. We previously used CRISPR/Cas9 to generate phospho-dead (S375A) and phospho-mimetic (S375E) alleles of *wipi-2* (Tsong *et al*., 2026). We found that neither *wipi-2* phosphomutant phenocopied *wipi-2(FTTG)* mutants in NSM dendrite length (Fig. S2A-D). While neither *wipi-2* phosphomutant displayed significantly different dorsal axon length from wild-type animals, *wipi-2(S375E)* mutants did have longer dorsal axons than *wipi-2(S375A)* mutant animals (Fig. S2E). Unsurprisingly, given that *wipi-2(null)* and *wipi-2(FTTG)* mutants did not display ectopic neurites, we also detected no significant effect of WIPI-2 phosphorylation on NSM ectopic neurites (Fig. S2F-G). In contrast, we did detect significant differences in NSM ventral axon arborization in the *wipi-2* phosphomutants, with *wipi-2(S375A)* phospho-dead mutants displaying higher ventral axon total length and more branchpoints than wild-type animals, while *wipi-2(S375E)* phospho-mimetic mutants were not different than wild-type animals (Figs. 4N, S2H-J). These results suggest that *wipi-2(S375A)* phospho-dead mutants have increased autophagosome biogenesis in the ventral axon and are consistent with dephosphorylated (phospho-dead) WIPI2 activating autophagy and with the *atg-13* rescuing lines similarly upregulating autophagy beyond normal levels (Fig. S1).

### ATG-9 is not necessary for restraining NSM ventral axon arborization

ATG-9 is a core autophagy protein across the eukaryotic evolutionary tree. To our knowledge, ATG9-independent autophagy has not been previously identified. Given our data indicating that *atg-9* is not necessary for NSM ventral axon arbor complexity (Fig. 4K-N), we first asked whether ATG-9 is present in the NSM ventral axon. We previously showed that ectopically expressed ATG-9 localizes to presynaptic regions in two other neurons in *C. elegans*, AIY and PVD. Further, we previously showed that endogenously labeled ATG-9 localizes similarly to RAB-3, a component of synaptic vesicles (Stavoe *et al*., 2016). Since the NSM ventral axon contains many RAB-3-positive presynaptic assemblies (Nelson and Colón-Ramos, 2013), we expected ATG-9 to also be present in the ventral axon, despite the lack of ventral axon arborization phenotype in *atg-9* mutants. Using the Native and Tissue-Specific Fluorescence (NATF) strategy (He *et al*., 2019), we endogenously labeled ATG-9 with three tandem copies of the eleventh β strand of GFP. We ectopically expressed GFP β strands 1-10 in NSM so that we could visualize endogenous ATG-9 localization only in NSM. Indeed, we observed ATG-9 present in the NSM ventral axon, including in ventral axon branches in L4 worms, indicating that endogenous ATG-9 is present in the ventral axon (Fig. 5A-B).

**Figure 5.**
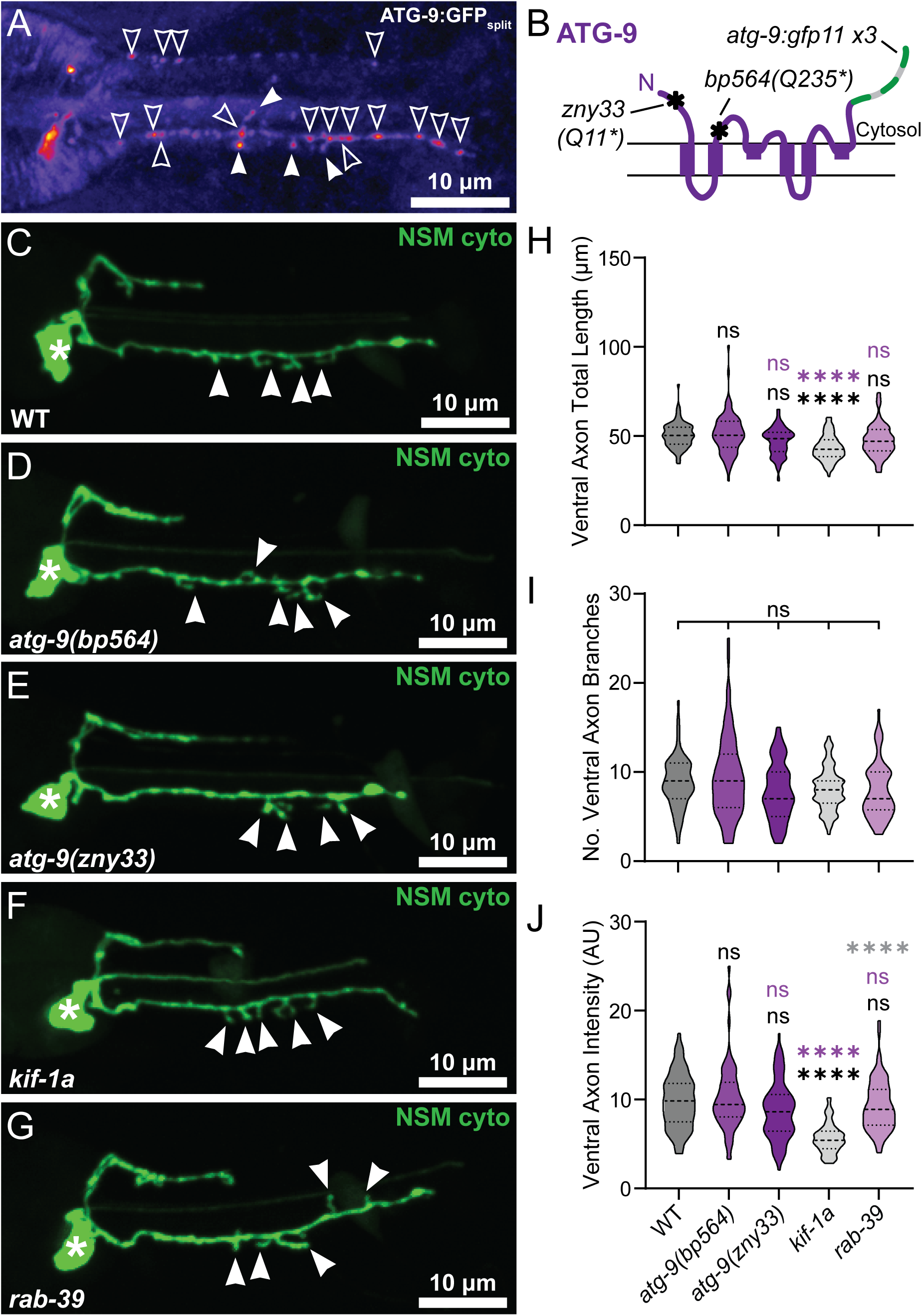
ATG-9 is not required to restrain NSM ventral axon arborization. (A) Representative maximal projection micrograph of NSM in L4 worms in a wild-type animal with endogenously tagged ATG-9 in NSM. Micrograph displayed with the Fire lookup table. Open arrowheads indicate ATG-9 signal in the dorsal and ventral axon shafts; closed arrowheads indicate ATG-9 signal in ventral axon branches. (B) Schematic of ATG-9 protein with location of alleles depicted. (C-J) Representative maximal projection micrographs of NSM in L4 worms in wild-type (C), *atg-9(bp564[Q235*])* (D), *atg-9(zny33[Q11*])* (E), *kif-1a/unc-104* (F), *rab-39* (G) mutant worms. Arrowheads indicate ventral axon branches; asterisk identifies the cell body. (H-J) Quantification of the total length of the ventral axon arbor, including branches (H), the number of ventral branchpoints (I), and the total intensity of the ventral axon arbor (J). Truncated violin plots with medium smoothing and median and quartiles displayed (n ≥ 70 cells imaged on at least three independent days for each genotype). ****p < 0.0001; ns > 0.05 between indicated groups, mutants and wild-type in black and between mutants, and *atg-9(bp564[Q235*])* in purple by Kruskal-Wallis test with Dunn’s Multiple Comparisons.

To confirm that ATG-9 does not phenocopy other autophagy mutants and is indeed dispensable for restraining NSM ventral axon arborization, we manipulated ATG-9 in the NSM axon in several ways (Fig. 5C-J). First, using CRISPR/Cas9, we generated an independent, novel allele of *atg-9* that introduces an early stop codon at ATG-9(Q11) (allele *zny33*) (Fig. 5B). The original *atg-9(bp564)* allele was generated using traditional mutagenesis methods and introduces an early stop codon at ATG-9(Q235) (Fig. 5B) (Tian *et al*., 2010). Our independent *atg-9(zny33)* allele behaved indistinguishably from the original *atg-9(bp564)* allele in NSM dendrite length and dorsal axon length (Fig. S3). Further, both *atg-9* alleles displayed congruent phenotypes in NSM ventral axon arbor total length, number of ventral axon branchpoints, and ventral axon total fluorescence (Fig. 5C-E, H-J). These data further indicate that while *atg-9* is required for restricting NSM dendrite outgrowth, *atg-9* is not necessary for NSM ventral axon arborization.

Next, we manipulated ATG-9 localization in the axon specifically. We previously found that UNC-104/KIF1A is required for ATG-9 axonal localization; in *kif-1a* mutants, ATG-9 is not present in the axon and is restricted to the cell body (Stavoe *et al*., 2016). Thus, in *kif-1a* mutants, we have effectively removed ATG-9 from the ventral axon, but maintained or increased its localization to the somatodendritic compartment. Consistent with our *atg-9* alleles, *kif-1a* mutants did not phenocopy other autophagy mutants in NSM ventral axon total length, number of ventral axon branches, or ventral axon total fluorescence (Fig. 5F, H-J). Indeed, *kif-1a* mutants had lower ventral axon total length (Fig. 5H) and ventral axon total intensity (Fig. 5J) than wild-type or *atg-9* mutant animals. KIF1A transports synaptic vesicles and other cargo into axons independent of ATG-9, which could explain the unique phenotypes for *kif-1a* mutants.

We also manipulated ATG-9 localization in the axon by using *rab-39* mutants. In Drosophila neurons, RAB-39 binds to KIF1A at the endosome in the cell body, removing a pool of KIF1A from transporting ATG-9 into the axon (Kilic *et al*., 2025). In *rab-39* mutants, this sequestration of KIF1A is released, allowing more ATG-9 to enter the axon. Similarly consistent with ATG-9 being dispensable for restraining NSM ventral axon complexity, we found that ventral axon total length, number of ventral axon branchpoints, and ventral axon total fluorescence were not different between wild-type and *rab-39* mutant animals (Fig. 5G-J).

To confirm the efficacy of the *kif-1a* and *rab-39* mutants, we quantified NSM dendrite length and dorsal axon length as well (Fig. S3). We expected *kif-1a* mutants to have the opposite phenotype of *atg-9* mutants in dendrite length, as a larger pool of ATG-9 would be available in the somatodendritic compartment in *kif-1a* mutants. Indeed, *kif-1a* mutants had shorter NSM dendrites than wild-type animals (Fig. S3F). In contrast, we observed that *rab-39* mutant worms exhibited NSM dendrites that were not significantly different from wild-type animals. The dorsal axon length was more variable in *kif-1a* mutant worms, similar to *atg-9* mutants (Fig. S3G). In contrast, *rab-39* mutant worms displayed significantly shorter dorsal axons than wild-type worms (Fig. S3G), which might be due to ATG-9-independent functions of RAB-39.

Taken together, our data indicate that ATG-9 is indeed not necessary to restrain NSM ventral axon arborization. We also observed that *atg-101* was not required to restrain ventral axon arborization (Fig. 4K-N). Interestingly, ATG101 is necessary to recruit ATG9 to the phagophore membrane (Ren *et al*., 2023), and ATG101 is required for ATG9 colocalization and interaction with ATG13 (Kannangara *et al*., 2021). Consequently, this role suggests that the absence of a ventral axon phenotype in *atg-101* mutants may be due to the lack of requirement of *atg-9* for NSM ventral axon arborization. Thus, we conclude that we have identified a novel, ATG9-independent form of autophagy.

### Autophagy-related scramblases can partially compensate for each other in NSM ventral axon arborization

ATG-9 is a lipid scramblase that moves a variety of lipids down their concentration gradients across the lipid bilayer at the growing phagophore membrane as they are delivered by the bridge-like lipid transfer protein ATG-2 (Holzer *et al*., 2024). Given the requirement for ATG-2 in restraining NSM ventral exon arborization, we hypothesized that some scramblase(s) should be required to move delivered lipids. Strikingly, ATG-9 has no related family members. In addition, scramblases cannot be bioinformatically identified based on sequence (Wang *et al*., 2022; Li *et al*., 2024; Rocha-Roa and Vanni, 2026). However, there are other scramblases that have been previously identified to operate in earlier steps related to autophagy. VMP1, TMEM41A and TMEM41B belong to the DedA family, an evolutionarily conserved family of proteins (Morita *et al*., 2018). VMP1 is a scramblase (Li *et al*., 2021) that resides in the ER and is necessary for autophagy in mammalian cells (Ropolo *et al*., 2007). Similarly, TMEM41B is a scramblase that is required for starvation-induced autophagy in HEK293T cells and physically and functionally interacts with VMP1 (Morita *et al*., 2018; Li *et al*., 2021). While both VMP1 and TMEM41B are indispensable for autophagy in mammalian immortalized cells, they also are known to participate in other molecular pathways. Worms have one VMP1 ortholog, EPG-3 (Tian *et al*., 2010), and one TMEM41B homolog, EGLI-1 (Da Silva *et al*., 2020). Worms also contain two proteins, BUS-19 and unnamed Y71A12C.2, that are orthologous to TMEM41A (Yook and Hodgkin, 2007; Kishore *et al*., 2024), a close relative of TMEM41B. We propose to name Y71A12C.2 “*tmem-41a.1*.”

We asked whether one or more of these scramblases were compensating for a loss of ATG-9 in NSM ventral axon arborization (Fig. 5A-M). Putative null alleles for *TMEM41B/egli-1*, *TMEM41A/bus-19* and *tmem-41a.1* were previously generated with traditional mutagenesis methods. We generated a putative null allele for *vmp-1* using CRISPR/Cas9, introducing an early stop codon (I95*). When we examined NSM ventral axon total length, number of ventral axon branchpoints, and ventral axon total fluorescence intensity in the scramblase mutants, we found no differences compared to wild-type animals (Fig. 6A-F, J-M), consistent with our findings in *atg-9* mutants and indicating that none of the autophagy-related scramblases are necessary to restrain NSM ventral axon arborization. We also quantified NSM dendrite length and dorsal axon length in the scramblase mutants (Fig. S4A-F). While *vmp-1* mutants displayed longer dendrites than wild-type animals, similar to autophagy mutants, the other scramblase mutants did not have longer NSM dendrites than wild-type worms (Fig. S4K). In addition, apart from *vmp-1* mutants, the other scramblase mutants showed consistently shorter NSM dorsal axons than both wild-type and *atg-9* mutant animals (Fig. S4K).

**Figure 6.**
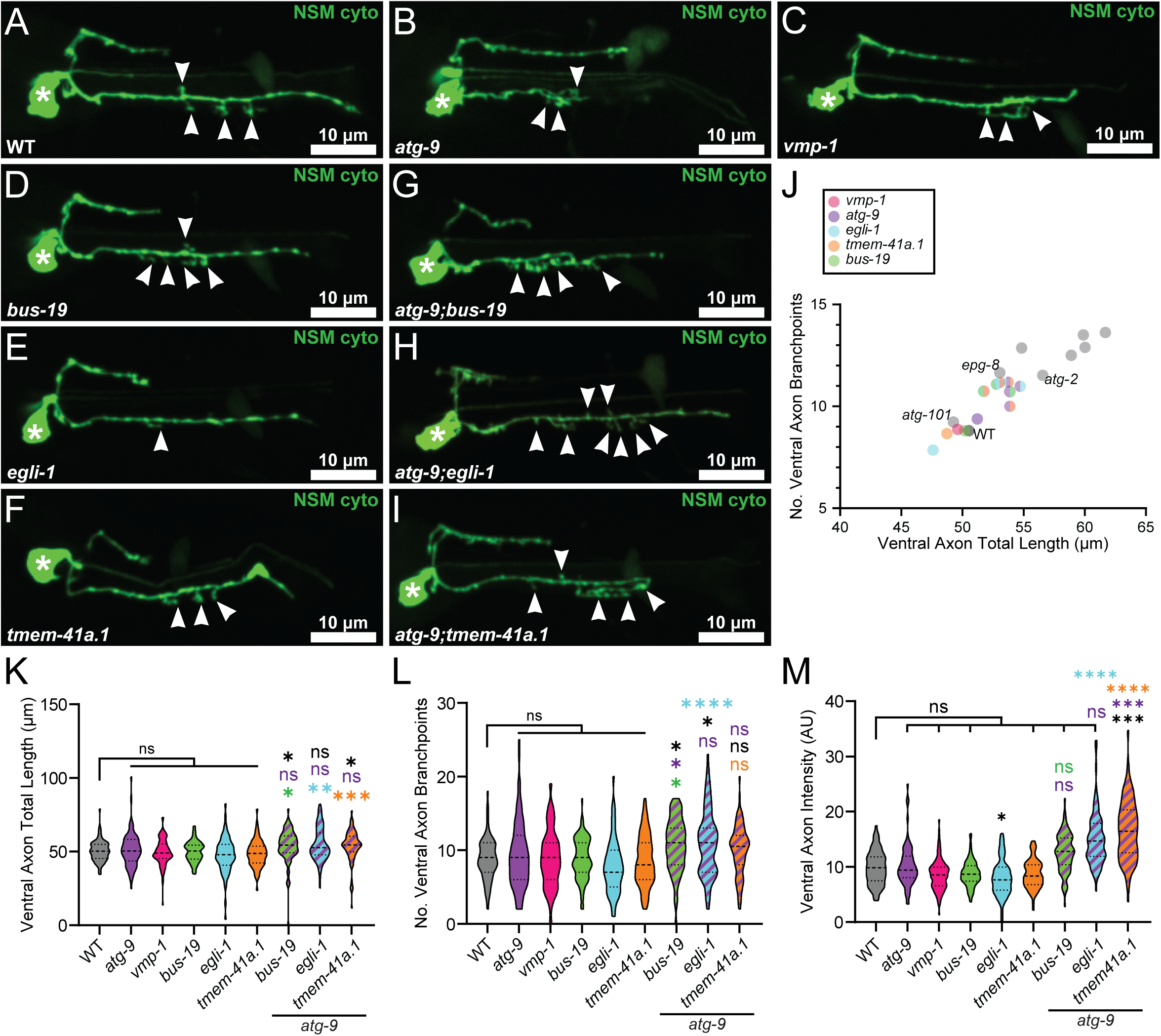
Scramblases can compensate for each in NSM ventral axon arborization. (A-I) Representative maximal projection micrographs of NSM in L4 worms in wild-type (A), *atg-9(bp564)* (B), *vmp-1/epg-3* (C), *bus-19* (D), *egli-1* (E), *tmem-41a.1* (F), *atg-9;bus-19* (G), *atg-9;egli-1* (H), and *atg-9;tmem-41a.1* (I) mutant worms. Arrowheads indicate ventral axon branches; asterisk identifies the cell body. (J) Plot of the means for number of branchpoints (y-axis) and ventral axon total length (x-axis) for indicated genotypes. Double and triple mutants are indicated by circles containing the relevant colors. (K-M) Quantification of the total length of the ventral axon arbor, including branches (K), quantification of the number of ventral branchpoints (L), and quantification of the total intensity of the ventral axon arbor (M). AU, arbitrary units. Truncated violin plots with medium smoothing and median and quartiles displayed (n ≥ 67 cells imaged on at least three independent days for each genotype). ****p < 0.0001; ***p < 0.0005; ** < 0.005; *p < 0.05; ns > 0.05 between mutants and wild-type in black and double mutants and their respective single mutants in the relevant colors by Kruskal-Wallis test with Dunn’s Multiple Comparisons.

To determine whether any of the scramblases can compensate for loss of *atg-9*, we generated double mutants between combinations of the scramblase mutants: *atg-9;bus-19* (Fig. 6G), *atg-9;egli-1* (Fig. 6H), *atg-9;tmem41a.1* (Fig. 6I), *bus-19;tmem41a.1* (Fig. S5A), *egli-1;tmem41a.1* (Fig S5C), and *bus-19;egli-1* (Fig. S5D). We also generated *atg-9;egli-1;tmem41a.1* triple mutants (Fig. S5B). We were not able to generate *atg-9;vmp-1* double mutants that were capable of producing viable progeny. We were interested to observe that the scramblase double mutants displayed some statistically significant phenotypes in NSM ventral axon arborization (Figs. 6J-M, S5E-G), particularly in NSM ventral axon branchpoints (Figs. 6L, S5F). Notably, the *atg-9;egli-1;tmem-41a.1* triple scramblase mutants displayed no more drastic NSM arborization phenotypes than the double scramblase mutants (Fig. S5). Thus, while the scramblase double mutants do show NSM arborization phenotypes, the NSM ventral axon phenotypes in scramblase double mutants are not as drastic as most of the autophagy mutants (Fig. 6J).

Moreover, the scramblase double mutants that contain atg-9(bp564) are not appreciably more drastic than those that do not (Fig. 6J), implying that ATG-9 does not occupy a unique scramblase role in NSM ventral axon branching. These data suggest that lipid scrambling is required for NSM ventral axon arborization but that autophagy-related scramblases can partially compensate for each other in restraining NSM ventral axon arborization.

Since autophagy-related scramblases can compensate for each other in NSM ventral axon arbor complexity (Fig. 6) and ATG-9 works in concert with the bridge-like lipid transfer protein (BLTP) ATG-2 (Holzer *et al*., 2024), we asked whether other BLTPs modulate NSM ventral axon arborization (Fig. S6A-H). BLTPs VPS-13A and BLTP-3B (Pandey *et al*., 2023) are highly expressed in NSM (Taylor *et al*., 2021; Liska *et al*., 2023). Yeast Vps13 works in parallel with Atg2 to transfer lipids to the growing phagophore (Dabrowski *et al*., 2023), and mammalian VPS13A can interact with ATG9 (van Vliet *et al*., 2024). We generated a putative null allele of *vps-13A(K194*)* using CRISPR/Cas9 and obtained a putative null allele of *bltp-3b*. We detected no significant differences in NSM ventral axon total length or number of branchpoints in either *vps-13a* or *bltp-3b* mutants compared to wild-type animals (Fig. S6A-C, S6F-G). Further, when we generated *atg-2;vps-13a* and *atg-2:bltp-3b* mutants, we observed no significant differences between the double mutants and *atg-2* single mutants in NSM ventral axon total length (Fig. S6F), number of branchpoints (Fig. S6G), or axon intensity (Fig. S6H). Together, these data suggest that ATG-2 is the sole BLTP that participates in autophagy-mediated regulation of NSM ventral axon arbor complexity. Taken together, our results indicate that multiple scramblases can compensate for each other to restrain NSM ventral axon arborization during development and suggest that the scramblases transfer lipids across the phagophore lipid bilayer after lipids are delivered to the phagophore by ATG-2.

### Autophagy promotes NSM ventral axon branch dynamicity during development

How does ATG-9-independent autophagy restrain NSM ventral axon arbor growth? To interrogate this question, we turned to time-lapse, live-animal imaging of NSM. We observed that NSM ventral axon branches grow and retract in L4 wild-type worms, with six branches retracting within a 35-minute imaging window (Fig. 7A, Movie 1). In contrast, NSM ventral axon branches do not retract as often in *atg-3* mutant L4 worms, with only one branch retracting within the same 35-minute imaging window (Fig. 7B, Movie 2). This is similar to autophagy restricting filopodia stability in Drosophila axons during development (Kiral *et al*., 2020). These data suggest that autophagy is required for NSM ventral axon branch retraction. When autophagy is lost, branches cannot retract, resulting in longer and more numerous axon branches in L4 animals.

**Figure 7.**
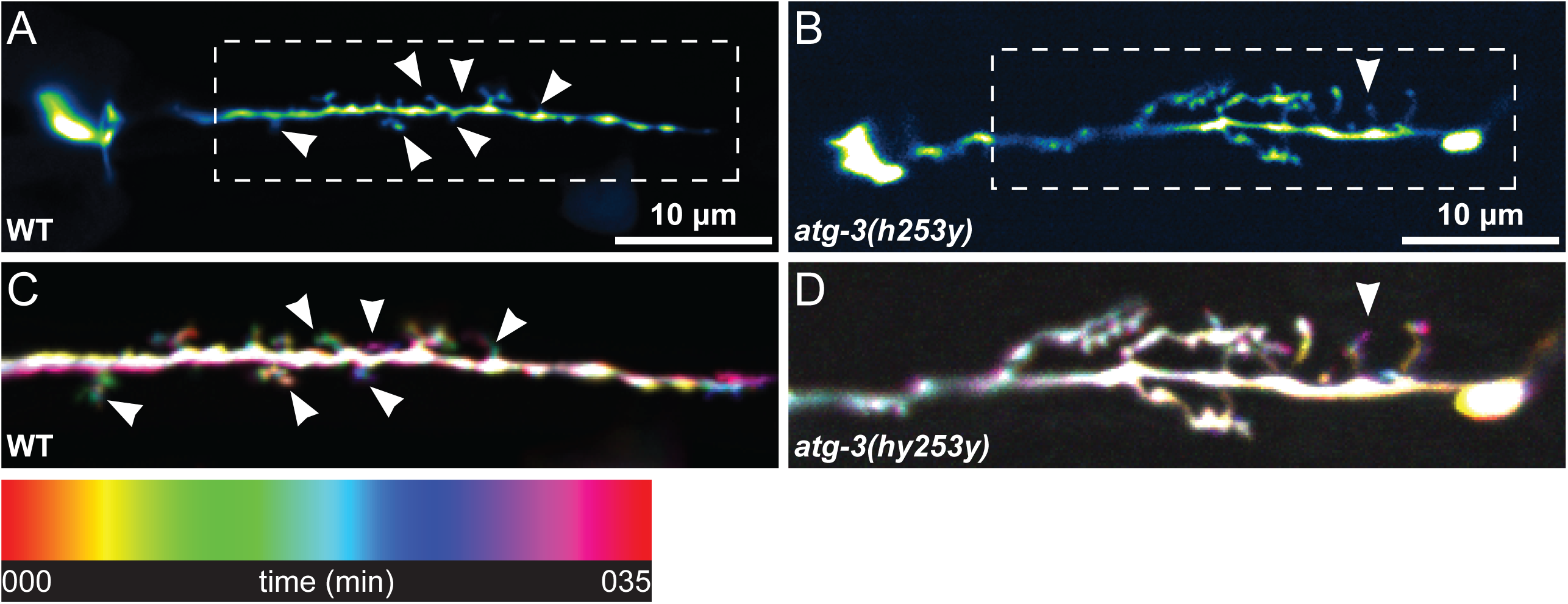
Autophagy promotes ventral axon branch dynamicity. (A-B) Still frames from live imaging of wild-type (A) and atg-3 mutant (B) animals. Arrowheads indicate dynamic branches and dashed boxes indicate areas of magnification below. Micrographs are displayed with the Green Fire Blue lookup table to more easily visualize the dim branches. (C-D) Magnifications indicated in A-B displayed with a temporal color code (legend below) using the Spectrum lookup table. Areas of the neurite present in all frames are white; colored branches are dynamic.

## DISCUSSION

We and others have previously shown that autophagy modulates discrete neurodevelopmental programs in individual neurons; for example, presynaptic assembly in interneuron AIY and axon outgrowth in sensory neuron PVD (Stavoe *et al*., 2016). Here, we determined that sensory/motor neuron NSM relies on autophagy to direct developmental programs in distinct compartments: soma, dendrite, dorsal axon, and ventral axon. NSM uses unique types of autophagy to restrict neurite outgrowth in each of its four compartments, underscoring the importance of the spatial regulation of autophagy in neurons. We have identified, to our knowledge, two novel forms of noncanonical autophagy: WIPI2-independent autophagy prevents ectopic neurite formation in the cell body and ATG9-independent autophagy restrains ventral axon arborization. Our findings in a single neuron shed light onto how a limited set of genes and molecular pathways can be shuffled to enable the formation of a highly complex, integrated nervous system. In addition, our data indicate that autophagy is broadly used in developing neurons to restrict neurite outgrowth in various neuronal compartments.

Several types of noncanonical autophagy have been previously identified, each of which is independent of at least one component of the canonical autophagy machinery (Dupont *et al*., 2021). ULK-independent autophagy can proceed in mammalian cell lines lacking ULK1 and ULK2 when cells are stressed with ammonia or hypoxia (Cheong *et al*., 2011; Feng *et al*., 2019). Similarly, Beclin1-independent autophagy responds to pro-death pathway inducers in yeast, *C. elegans*, and mammalian cells (Zhu *et al*., 2007; Scarlatti *et al*., 2008; Grishchuk *et al*., 2011; Mauthe *et al*., 2011). However, PI3P generation still appears to be required for autophagosome biogenesis, as Class II phosphatidylinositol 3-kinase (PIK3C2) can generate PI3P instead (Devereaux *et al*., 2013; Boukhalfa *et al*., 2020). Alternatively, it has been proposed that PI5P can substitute for PI3P in the nucleation of phagophores during autophagosome biogenesis (Vicinanza *et al*., 2015; Nascimbeni *et al*., 2017). Additionally, conjugation-independent autophagy can generate double-membrane vesicles identifiable by ultrastructural studies in mammalian cells lacking all six mammalian ATG8s (Nguyen *et al*., 2016) or in cells lacking ATG3, ATG5, or ATG7 (Mizushima *et al*., 2001; Sou *et al*., 2008; Nishida *et al*., 2009; Kishi-Itakura *et al*., 2014; Uemura *et al*., 2014).

In our systematic assessment of the role of autophagy in NSM development, we have identified two novel forms of noncanonical autophagy. WIPI2-independent autophagy blocks ectopic neurite outgrowth in the NSM cell body (Fig. 3). This form of autophagy appears to require PI3P, as both ATG-14, a component of the PI3P-generating nucleation complex, and WIPI-4, a PROPPIN that binds to PI3P, are required to prevent ectopic neurites. Further, in canonical autophagy, WIPI2 bridges the nucleation and elongation complexes by binding PI3P and ATG16L1 (Dooley *et al*., 2014). However, the elongation complex is also required for curbing the development of ectopic neurites, as *atg-3* mutants display a strong ectopic neurite phenotype. Therefore, it is possible that another protein can compensate for the loss of WIPI-2 or that the elongation complex is recruited to the PI3P-decorated phagophore by a different mechanism.

We were similarly surprised to identify ATG-9-independent autophagy controlling ventral axon arborization (Figs. 4, 5). Conjugation-independent autophagy (also called alternative autophagy) is also independent of ATG9 (Tsuboyama *et al*., 2016). Similarly, PIK3C3-indepdent autophagy does not require ATG9 (Devereaux *et al*., 2013; Dupont *et al*., 2021). However, the conjugation machinery and nucleation complex are both necessary for restricting NSM ventral axon arborization (Fig. 4). The requirement for BLTP ATG-2 for ventral arborization suggests that lipids delivered to the cytoplasmic phagophore leaflet by ATG-2 would need to be moved to the luminal leaflet by a scramblase to relieve physical tension in the phagophore membrane. The other autophagy-associated scramblases, VMP-1, TMEM-41B and its relative TMEM-41A, are not known to localize to the phagophore, but instead to the ER (Moretti *et al*., 2018; Morita *et al*., 2018; Shoemaker *et al*., 2019). Further, we know of no studies that indicate VMP-1, TMEM-41B or TMEM-41A can compensate for the loss of ATG-9 in *C. elegans* or other systems. However, while autophagosome numbers decreased significantly in Atg9 knockout mouse embryonic fibroblasts, some LC3B^+^ puncta were still observed (Saitoh *et al*., 2009), suggesting that conserved compensatory pathways may exist to enable limited autophagosome formation. Additionally, our inability to generate viable *atg-9;vmp-1* double mutants provides additional evidence that VMP-1 and ATG-9 act redundantly in some aspect(s) of reproduction. It is intriguing that loss of any two of the scramblases ATG-9, TMEM-41A (BUS-19 and TMEM-41A.1), and TMEM-41B (EGLI-1) was sufficient to generate a moderate phenotype in NSM ventral axon arborization, suggesting that the identity of the scramblase is not particularly important for ventral axon arborization. Given the lack of sequence similarity between ATG-9 and the other scramblases (Wang *et al*., 2022; Li *et al*., 2024; Rocha-Roa and Vanni, 2026), it is unclear how other scramblases could be molecularly recruited to the phagophore to scramble lipids as necessary. However, there are some intriguing hints from other studies that it may be possible. First, ATG9 can colocalize with TMEM41B in mammalian cells (Mailler *et al*., 2021). In addition, while the ATG2 N-terminus can associate with TMEM41B and VMP1 (Ghanbarpour *et al*., 2021), ATG9 has been shown to be able to interact with either the C-or N-terminus of ATG2 (Wang *et al*., 2024).

Despite the compartment specificity and apparent disparity between the NSM phenotypes we identified, the phenotypes appear to converge on neurite outgrowth. NSM dendrite length at L4 (Fig. 1) can be the result of increased neurite outgrowth in autophagy mutants. Similarly, dorsal axon length at L4 is a direct consequence of neurite outgrowth (Fig. 2), although the role of autophagy is less clear. The NSM ectopic neurites we observed in autophagy mutants other than *wipi-2* mutants (Fig. 3) are more likely the result of derepression of initiation of neurite outgrowth. Finally, while we did not observe differences in the length of the main NSM ventral axon (data not shown), the increase in the number of axon branches and ventral axon total length in autophagy mutants other than *atg-9* mutants (Fig. 4) is likely the result of lower rates of branch collapse or retraction (Fig. 7). Similarly, we previously showed that autophagy mutants display longer PVD axons at L4 (Stavoe *et al*., 2016) and others have demonstrated that inhibition of autophagy induces axon outgrowth in primary mammalian neurons (Tamura and Ohkuma, 1991; Ban *et al*., 2013). Altogether, these data suggest that autophagy may control neurodevelopment by broadly restricting neurite outgrowth, possibly via the RhoA-ROCK signaling pathway as seen in primary mammalian cortical neurons (Ban *et al*., 2013). By restraining neurite outgrowth across neurons, autophagy may enable neurons to find their proper synaptic partners and develop the correct neural circuits that ultimately underlie behaviors, memory and cognition.

## METHODS

### Reagents

#### Strains

We maintained *C. elegans* strains at room temperature on NGM plates seeded with OP50 *E. coli*. We obtained strain LX837 *vsIs45* from the *Caenorhabditis* Genetics Center (CGC) and used it as our wild-type control strain in this study. Some strains were provided by the CGC, which is funded by NIH Office of Research Infrastructure Programs (P40 OD010440): CB1265 *unc-104(e1265)*, CB6598 *bus-19(e2966)*, HZ1683 *atg-2(bp576);him-5(e1490)*, HZ1687 *atg-9(bp564)*, HZ1688 *atg-13(bp414)*, HZ1690 *epg-6(bp424);him-5(e1490)*, HZ1692 *epg-9(bp320);him-5(e1490)*, PS9934 *rab-39(sy1987)*, VC577 *egli-1(gk278)*, VC30247 *atg-18(gk447069)*, VC40643 *tmem-41a.1/y71A12c.2(gk940818)*, and VC40484 *bltp-3b(gk660281)*. Information about the specific alleles can be found in Supplementary Table 1. Any strains that we received from the CGC were genotyped upon receipt by PCR and restriction digest (when applicable).

#### Plasmids

We generated tph-1p::atg-13 cDNA::SL2::TagBFP (pwAS386) in the pSM backbone with restriction enzyme subcloning from existing plasmids. The plasmid contains the rab-3 3’UTR.

#### CRISPR Transgenics

We adapted and used a co-CRISPR protocol (Ghanta and Mello, 2020) to generate *atg-9(zny33[Q11*])*, *epg-3(zny36[I95*])*, *epg-8(zny40[V238*])*, and *vps-13a(zny39[K194*])*. We recapitulated the *dpy-10(cn64)* lesion to generate roller and dumpy worms as the co-CRISPR marker (Dickinson and Goldstein, 2016). We designed all CRISPR strategies and obtained all CRISPR reagents from Integrated DNA Technologies (IDT). We chose guide RNAs and lesions to reduce or eliminate predicted off-target hits. We optimized the *dpy-10(cn64)* guide RNA sequence to eliminate off-target hits (sequence: CATAGGCACCACGAGCGGTA). We performed CRISPR in LX837 *vsIs45* worms and confirmed CRISPR edits with Sanger sequencing. For *atg-9(ola274[atg-9::gfp11x3])*, we inserted the sequence “ggaggggcatccgcagggggaggtcgcgatcacatggtcctgcatgagtatgtgaacgccgccgggatcactggtggctctggaggtagagatcatatggttctccacgaatacgttaacgccgcaggcatcactggcggtagtggaggacgcgaccatatggtactacatgaatatgtcaatgcagccggaataacc” immediately in front of the TAG stop codon in *atg-9*. For *atg-3(ola502)*, we deleted the sequence between aatttaaaaagtgct and atactcgtgctattaaattgt. We confirmed that no off-target lesions (as predicted by IDT CRISPR design tool) were present in the background by Sanger sequencing or whole-genome sequencing (University of Minnesota Model Organism Sequencing Service).

#### Microscopy

We mounted worms onto 2% agarose pads and immobilized worms with 10 mM levamisole (MilliporeSigma) prior to imaging. We acquired z-stacks (0.3 μm step size) on a spinning-disk confocal microscope (Nikon Ti2 Inverted Confocal with Yokogawa W1 Spinning Disk Package) with an Apochromat 60x, 1.4 NA oil immersion objective (Nikon Instruments) with a back-illuminated cCMOS camera (Teledyne Photometrics) using Nikon Elements software.

#### Time-lapse, live-animal microscopy

We immobilized L4 stage animals on 10% agarose pads with 10mM levamisole. We used a Nikon Ti2 microscope equipped with a CSU-W1 spinning disk head, ORCA-Fusion BT SCMOS camera, high-speed piezo stage motor, 60X, 1.40 NA Apo Lambda oil objective lens for time-lapse imaging. We acquired single-plane images every 30 or 60 seconds for more than 25 minutes to follow dynamics of axonal branches. We used the ND alignment function on NIS Elements AR analysis software (version 6.10.01) to align and correct for image drift on the whole image and produce supplementary videos. We cropped and annotated videos in FIJI (Image J).

#### Micrograph Analysis and Quantification

We generated maximal intensity projections of z-stacks in Nikon Elements. We measured dendrite length, dorsal axon length, and soma ectopic neurites on maximal intensity projection micrographs in FIJI (Schindelin *et al*., 2012). We confirmed the distal end of neurites by referencing the original z-stacks when needed.

To quantify the ventral axon arbor complexity, we first isolated the ventral axon from the maximal intensity projection micrograph in FIJI. We then developed an unbiased analysis pipeline in Nikon Elements using General Analysis 3 (GA3). In the GA3, we had Elements threshold the image and then identify and segment the axon.

#### Statistics

We assembled data in Microsoft Excel and GraphPad Prism 11. We generated graphs and performed statistical tests in Prism 11. We tested data for normality with the D’Agostino and Pearson test. Based on this analysis of normality, we subsequently used the appropriate statistical test, which are indicated in the corresponding figure legends.

## Key Resources Table

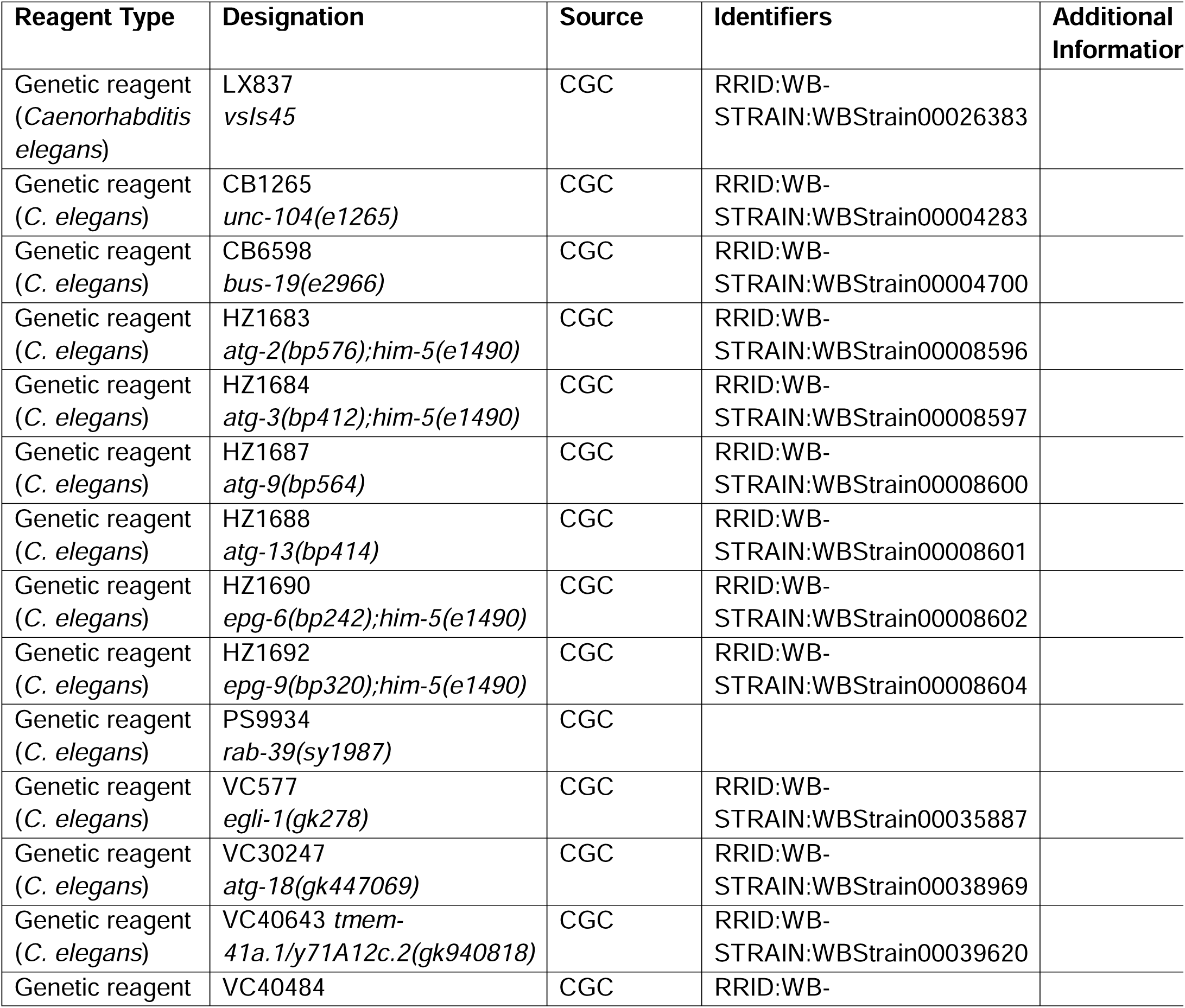
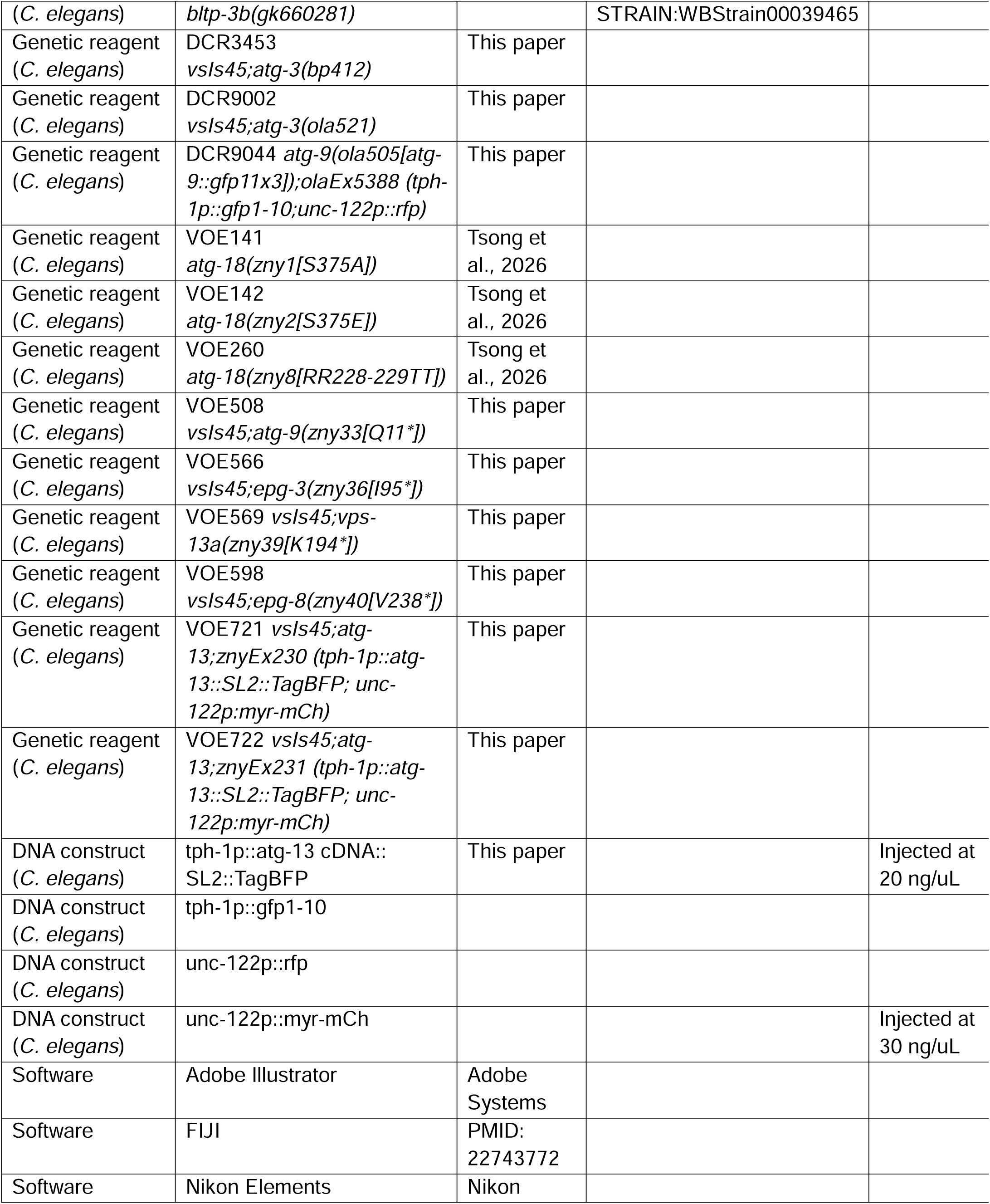
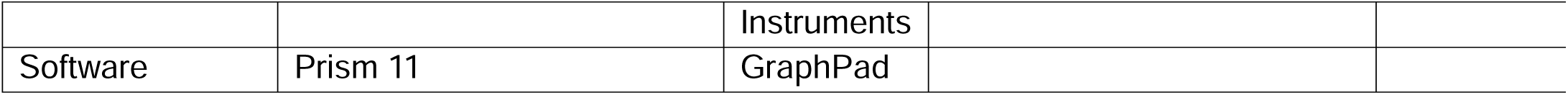

## ACKNOWLEDGEMENTS

The authors acknowledge the Waxham and Arey labs for helpful discussions and Grace Augustine and Beverly Hughes for technical assistance.

## SUPPLEMENTAL FIGURE LEGENDS

**Figure S1. Autophagy acts cell-autonomously in NSM.**

(A-C) Representative maximal projection micrographs of NSM in L4 worms in *atg-13* mutants (A) and *atg-13* mutants with rescuing line 1 (B) and rescuing line 2 (C). Closed arrowheads indicate ventral axon branches; open arrowhead indicates the distal end of the dorsal axon; arrow indicates the distal end of the dendrite; asterisk identifies the cell body. Bottom micrographs are displayed with the Fire lookup table to more easily visualize the dim NSM dendrite. (D-J) Quantification of NSM neurodevelopment. (D) Quantification of dendrite length. Truncated violin plot with medium smoothing and median and quartiles displayed (n ≥ 152 cells imaged on at least three independent days for each genotype). (E) Quantification of the penetrance of NSM ectopic neurites in wild-type and autophagy mutant animals. Error bars indicate 95% confidence interval; n > 151 cells imaged on at least three independent days for each genotype. ****p < 0.0001 between autophagy mutants and wild-type in black and between rescuing lines and *atg-13* mutants in dark red; ns > 0.05 between rescuing lines by Fisher’s exact test. (F-I) Truncated violin plot with medium smoothing and median and quartiles displayed. (F) Quantification of the number of ectopic neurites per cell (n ≥ 151 cells imaged on at least three independent days for each genotype). (G) Quantification of the total length of ectopic neurites per cell (n ≥ 152 cells imaged on at least three independent days for each genotype). (H) Quantification of the total length of the ventral axon arbor, including branches (n ≥ 76 cells imaged on at least three independent days for each genotype). (I) Quantification of the number of ventral branchpoints (n ≥ 76 cells imaged on at least three independent days for each genotype). For F-I, ****p < 0.0001; ***p < 0.0005; **p < 0.005; *p < 0.05; ns > 0.05 between indicated groups or between mutants and wild-type in black and between rescuing lines and *atg-13* mutants in dark red by Kruskal-Wallis test with Dunn’s Multiple Comparisons. (J) Plot of the means for number of branchpoints (y-axis) and ventral axon total length (x-axis) for indicated genotypes. Unlabeled gray points correlate with labeled autophagy mutants in Fig. 4N for reference.

**Figure S2. WIPI-2 phosphorylation in NSM neurodevelopment.**

(A-C) Representative maximal projection micrographs of NSM in L4 worms in wild-type animals (A), *wipi-2/atg-18(S375A)* (B), and *wipi-2/atg-18(S375E)* (C) animals. Closed arrowheads indicate ventral axon branches; open arrowhead indicates the distal end of the dorsal axon; arrow indicates the distal end of the dendrite; asterisk identifies the cell body. Bottom micrographs are displayed with the Fire lookup table to more easily visualize the dim NSM dendrite. (D-J) Quantification of NSM neurodevelopment. Truncated violin plot with medium smoothing and median and quartiles displayed. (D) Quantification of dendrite length (n ≥ 139 cells imaged on at least three independent days for each genotype). (E) Quantification of dorsal axon length (n ≥ 85 cells imaged on at least three independent days for each genotype). (F) Quantification of the number of ectopic neurites per cell (n ≥ 149 cells imaged on at least three independent days for each genotype). (G) Quantification of the total length of ectopic neurites per cell (n ≥ 146 cells imaged on at least three independent days for each genotype). (H) Quantification of the total length of the ventral axon arbor, including branches (n ≥ 73 cells imaged on at least three independent days for each genotype). (I) Quantification of the number of ventral branchpoints (n ≥ 73 cells imaged on at least three independent days for each genotype). (J) Quantification of the total intensity of the ventral axon arbor (n ≥ 72 cells imaged on at least three independent days for each genotype). AU, arbitrary units. ****p < 0.0001; ***p < 0.0005; ** < 0.005; *p < 0.05; ns > 0.05 between mutants and wild-type in black, between mutants and *wipi-2(FTTG)* in teal, and between *wipi-2* phosphomutants in berry by Kruskal-Wallis test with Dunn’s Multiple Comparisons.

**Figure S3. Manipulating ATG-9 in the NSM axon in NSM dendrite and dorsal axon outgrowth.**

(A-E) Representative maximal projection micrographs of NSM in L4 worms in wild-type (A), *atg-9(bp564[Q235*])* (B), *atg-9(zny33[Q11*])* (C), *kif-1a/unc-104* (D), *rab-39* (E) mutant worms. Open arrowhead indicates the distal end of the dorsal axon; arrow indicates distal end of the dendrite; asterisk identifies the cell body. (F) Quantification of dendrite length. Truncated violin plot with medium smoothing and median and quartiles displayed (n ≥ 139 cells imaged on at least three independent days for each genotype). (G) Quantification of dorsal axon length. Truncated violin plot with medium smoothing and median and quartiles displayed (n ≥ 133 cells imaged on at least three independent days for each genotype). For F-G, ****p < 0.0001; ***p < 0.0005; ns > 0.05 between mutants and wild-type in black, between mutants and *atg-9(bp564[Q235*])* in purple, and between *rab-39* and *kif-1a/unc-104* mutans in gray by Kruskal-Wallis test with Dunn’s Multiple Comparisons.

**Figure S4. The role of scramblases in NSM dendrite and dorsal axon outgrowth.**

(A-I) Representative maximal projection micrographs of NSM in L4 worms in wild-type (A), *bus-19* (B), *egli-1* (C), *tmem-41a.1* (D), *vmp-1/epg-3* (E), *atg-9(bp564)* (F), *atg-9;bus-19* (G), *atg-9;egli-1* (H), and *atg-9;tmem-41a.1* (I) mutant worms. Open arrowhead indicates the distal end of the dorsal axon; arrow indicates the distal end of the dendrite; asterisk identifies the cell body. (J) Quantification of dendrite length. Truncated violin plot with medium smoothing and median and quartiles displayed (n ≥ 148 cells imaged on at least three independent days for each genotype). (K) Quantification of dorsal axon length. Truncated violin plot with medium smoothing and median and quartiles displayed (n ≥ 144 cells imaged on at least three independent days for each genotype). For J and K (legend above J), ****p < 0.0001; *p < 0.05; ns > 0.05 between indicated groups or to wild-type in black; to *atg-9(bp564)* in purple; # indicates significance to both wild-type and *atg-9(bp564)*; $ indicates ****p < 0.0001 to wild-type and ns > 0.05 to *atg-9(bp564)*; & indicates ns > 0.05 to wild-type and ****p < 0.0001 to *atg-9(bp564)* by Kruskal-Wallis test with Dunn’s Multiple Comparisons.

**Figure S5. Scramblases in NSM ventral axon arborization.**

(A-D) Representative maximal projection micrographs of NSM in L4 worms in *bus-19;tmem-41a.1* (A), *atg-9;egli-1;tmem-41a.1* (B), *egli-1;tmem-41a.1* (C), and *bus-19;egli-1* (D) mutant worms. (E-G) Quantification of the total length of the ventral axon arbor, including branches (E), quantification of the number of ventral branchpoints (F), and quantification of the total intensity of the ventral axon arbor. AU, arbitrary units (G). Truncated violin plots with medium smoothing and median and quartiles displayed (n ≥ 67 cells imaged on at least three independent days for each genotype). ****p < 0.0001; ***p < 0.0005; ** < 0.005; *p < 0.05; ns > 0.05 between mutants and wild-type in black and double mutants and their respective single mutants in the relevant colors by Kruskal-Wallis test with Dunn’s Multiple Comparisons.

**Figure S6. BLTPs in NSM ventral axon arborization.**

(A-E) Representative maximal projection micrographs of NSM in L4 worms in *atg-2* (A), *vsp-13a* (B), *bltp-3b* (C), *vps-13a;atg-2* (D) and *bltp-3b;atg-2* (E) mutant worms. (F-H) Quantification of the total length of the ventral axon arbor, including branches (F), quantification of the number of ventral branchpoints (G), and quantification of the total intensity of the ventral axon arbor. AU, arbitrary units (H). Truncated violin plots with medium smoothing and median and quartiles displayed (n ≥ 69 cells imaged on at least three independent days for each genotype). ****p < 0.0001; ***p < 0.0005; ** < 0.005; *p < 0.05; ns > 0.05 between mutants and wild-type in black and double mutants and their respective single mutants in the relevant colors by Kruskal-Wallis test with Dunn’s Multiple Comparisons.

**Table S1. Gene orthologs and alleles.**

Gene names used in the paper are listed with their corresponding official worm and mammalian orthologs. Alleles used in this paper are delineated with the associated genetic sequences for the wild-type and mutant alleles. Autophagy genes are listed first and are grouped by autophagy complex.

**Movie 1. NSM ventral branch dynamics in a wild-type animal.**

Time-lapse movie of the ventral axon arbor of NSM in a L4 wild-type animal. Arrows indicate branches that display dynamicity during the imaging window. Axon depicted with Green Fire Blue lookup table. Scale bar, 10 µm. Still frames and color time scale depicted in Figure 7.

**Movie 2. NSM ventral branch dynamics in an *atg-3* mutant animal.**

Time-lapse movie of the ventral axon arbor of NSM in a L4 *atg-3* mutant animal. Arrow indicates branch that displays dynamicity during the imaging window. Axon depicted with Green Fire Blue lookup table. Scale bar, 10 µm. Still frames and color time scale depicted in Figure 7.

