## Supplementary figures and images for "Different forms of autophagy restrict neurite outgrowth in disparate compartments in a single neuron"

### Supplemental Figures

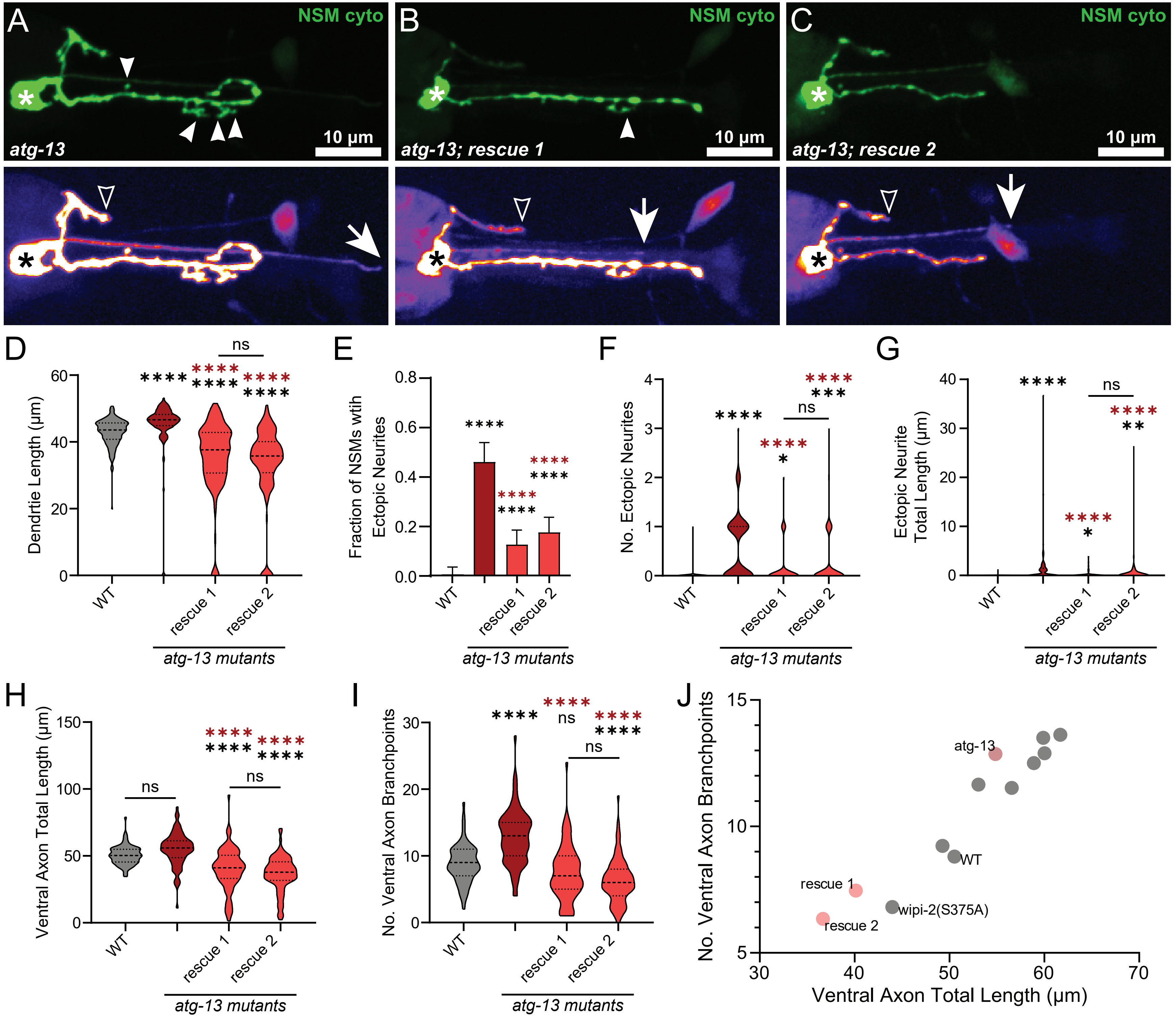

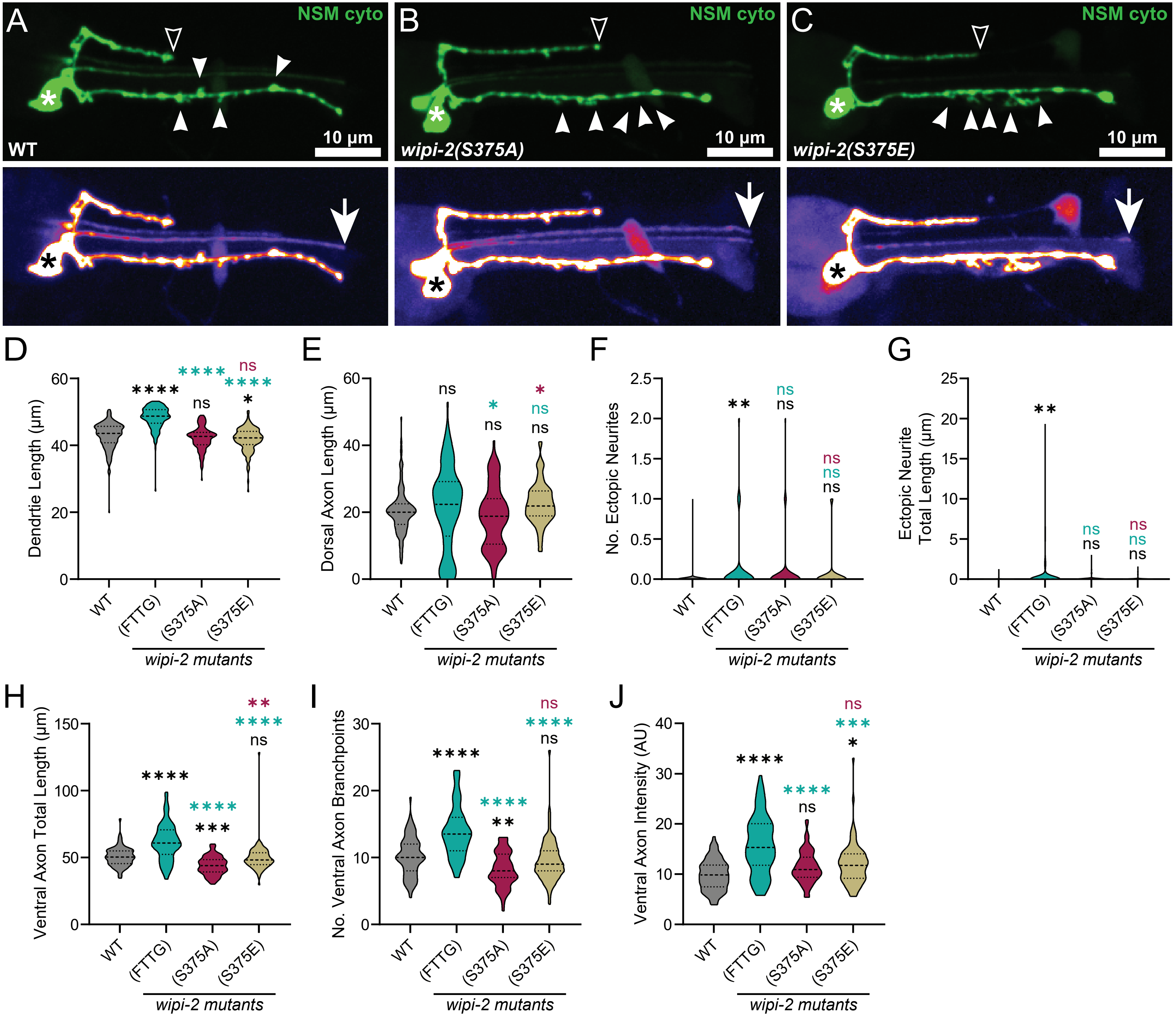

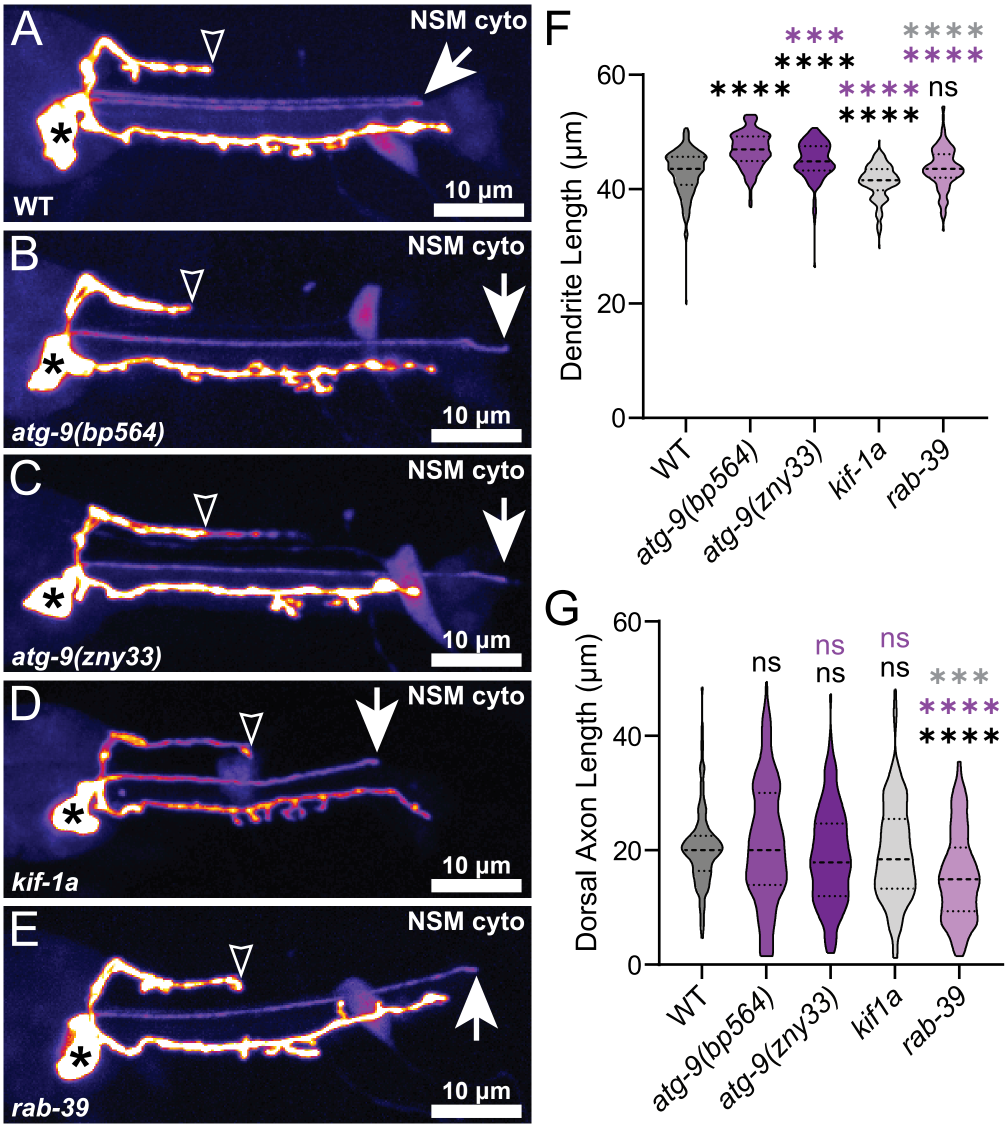

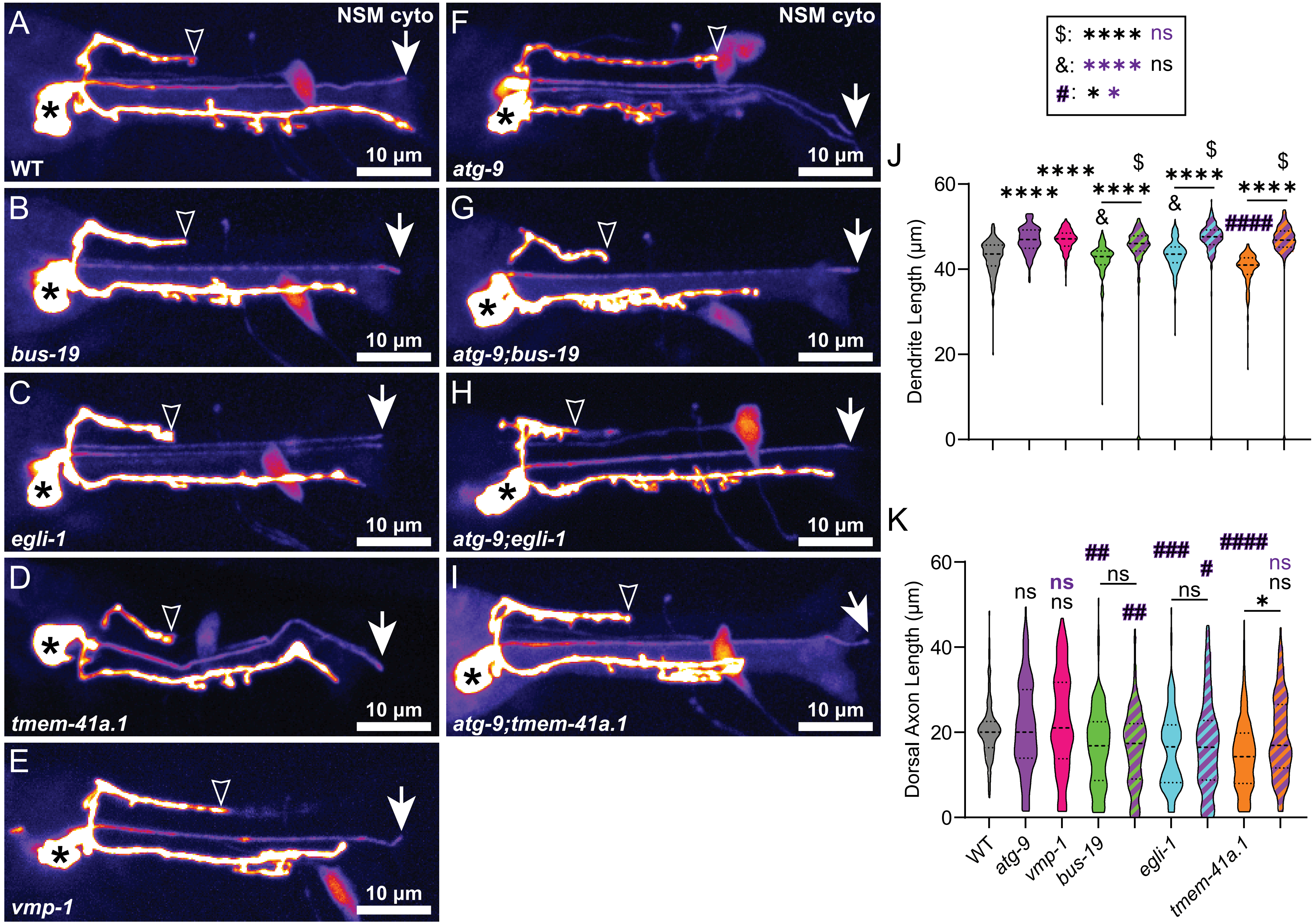

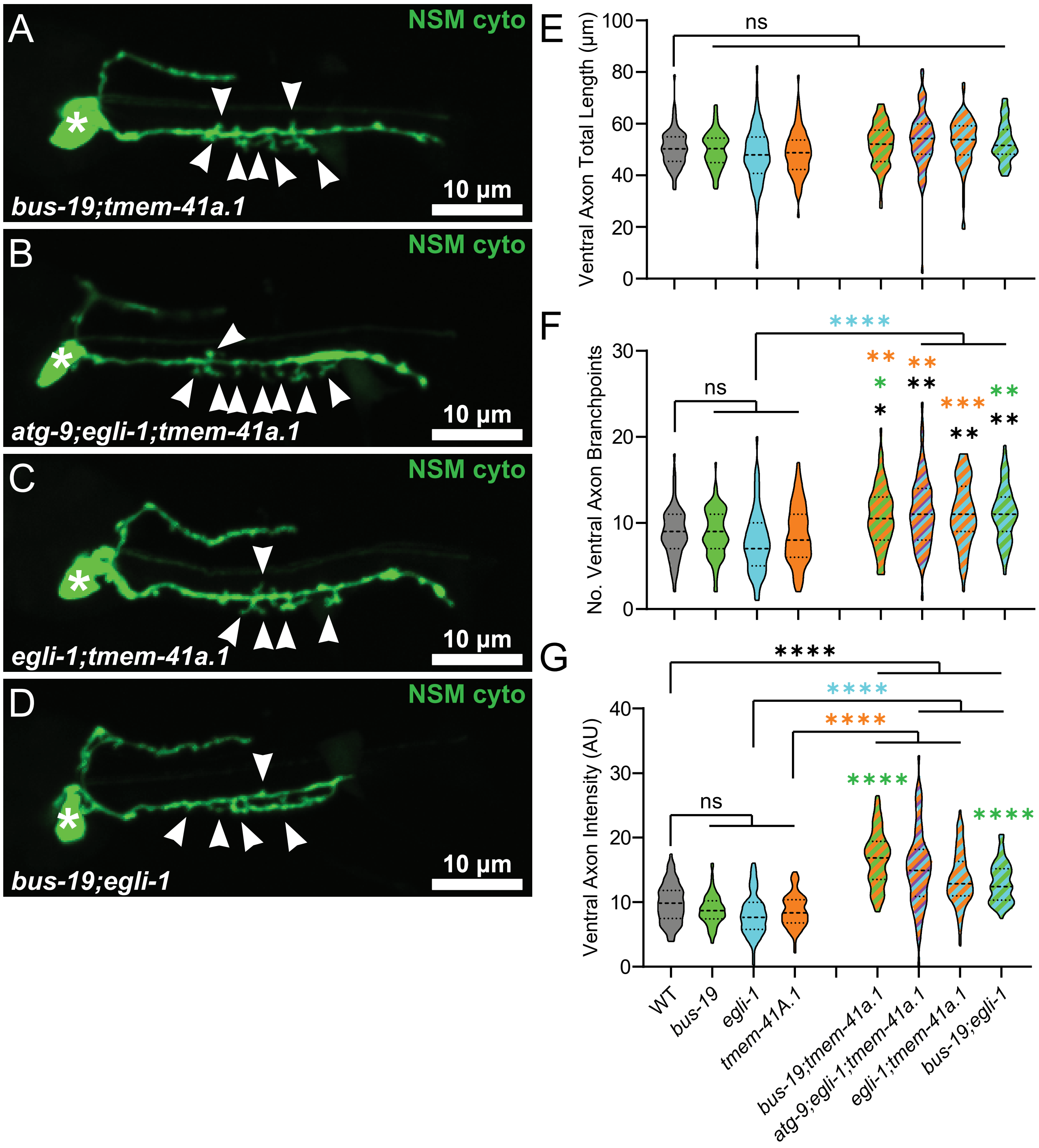

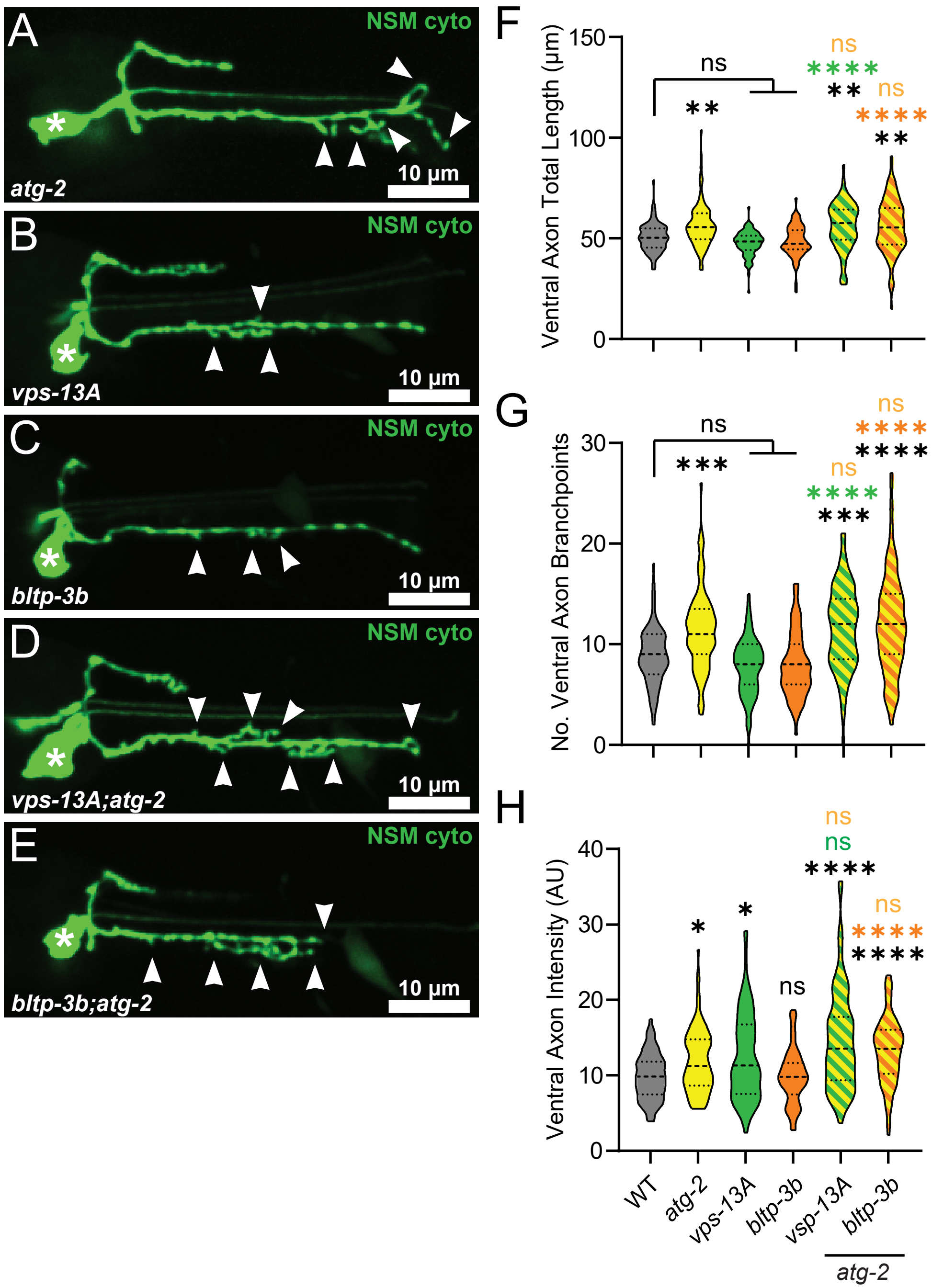
